# Extended hopanoid loss reduces bacterial motility and surface attachment and leads to heterogeneity in root nodule growth kinetics in a *Bradyrhizobium-Aeschynomene* symbiosis

**DOI:** 10.1101/423301

**Authors:** B.J. Belin, E.T Tookmanian, J. de Anda, G. C. L Wong, D.K. Newman

## Abstract

Hopanoids are steroid-like bacterial lipids that enhance membrane rigidity and promote bacterial growth under diverse stresses. Hopanoid biosynthesis genes are conserved in nitrogen-fixing plant symbionts, and we previously found that the extended (C_35_) class of hopanoids in *Bradyrhizobium diazoefficiens* are required for efficient symbiotic nitrogen fixation in the tropical legume host *Aeschynomene afraspera*. Here we demonstrate that the nitrogen fixation defect conferred by extended loss can fully be explained by a reduction in root nodule sizes rather than per-bacteroid nitrogen fixation levels. Using a single-nodule tracking approach to track *A. afraspera* nodule development, we provide a quantitative model of root nodule development in this host, uncovering both the baseline growth parameters for wild-type nodules and a surprising heterogeneity of extended hopanoid mutant developmental phenotypes. These phenotypes include a delay in root nodule initiation and presence of a subpopulation of nodules with slow growth rates and low final volumes, which are correlated with reduced motility and surface attachment *in vitro* and lower bacteroid densities *in planta*, respectively. This work provides a quantitative reference point for understanding the phenotypic diversity of ineffective symbionts in *A. afraspera* and identifies specific developmental stages affected by extended hopanoid loss for future mechanistic work.

## Introduction

Hopanoids are steroid-like lipids that support bacterial survival under stress (reviewed in Belin et al. 2018). They are synthesized by the squalene-hopene cyclase (*shc*) family of enzymes (Ochs et al. 1992; Syren et al. 2016), which generate the pentacyclic, C_30_ hopanoid core from squalene. In many organisms, the C_30_ hopanoids can be further modified, including methylation at the C-2 position via the enzyme HpnP (Welander et al. 2010) and addition of a ribose-derived side chain by the enzyme HpnH (Fig. 1a) (Welander et al. 2012). Side chain-containing hopanoids are known collectively as the C_35_ or “extended” hopanoids and commonly include molecules with aminotriol-, polyol-, and adenosyl-side-chain moieties (Schmerk et al. 2015). Organism-specific side chains have also been observed, including a hopanoid-lipid A conjugate known as HoLA (Silipo et al. 2014; Kulkarni et al. 2015; Komaniecka et al. 2014) that so far has only been found in *Bradyrhizobiaceae*.

**Figure 1.**
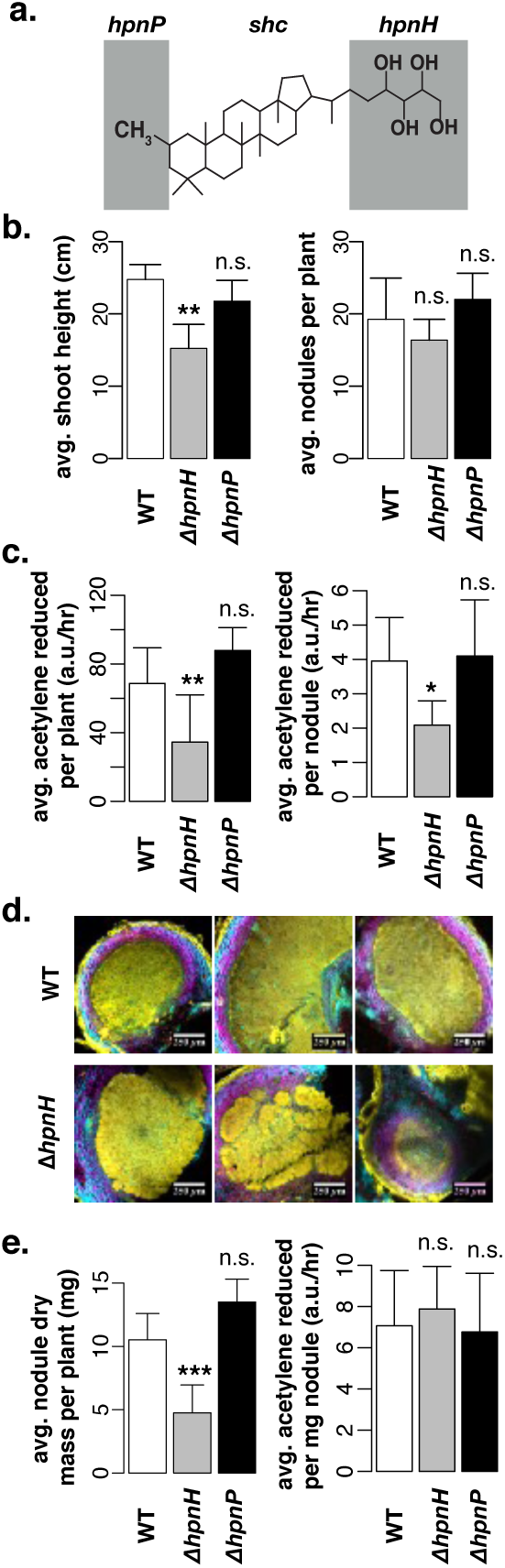
The nitrogen fixation defect of Δ*hpnH* results from a reduction in nodule sizes. (**a**) Chemical structure of the extended hopanoid 2-Methyl Bacteriohopanetetrol (2Me-BHT), consisting of a central pentacyclic core synthesized by the *shc* gene product, a C2 methylation site added by the product of *hpnP* (grey shading, left), and a tetrol group added by the *hpnH* product (grey shading, right). (**b**) Average shoot heights and nodules per plant at 24 dpi for *A. afraspera* plants inoculated with wild-type, Δ*hpnH* or Δ*hpnP B. diazoefficiens.* (**c**) Average acetylene reduction per plant and per nodule at 24 dpi for *A. asfrapera* plants inoculated with wild-type, Δ*hpnH* or Δ*hpnP*. (**d**) Representative confocal images of cross-sections of wild type- and Δ*hpnH*-infected nodules at 24 dpi illustrating plant cell walls (Calcofluor, cyan), live bacteria (SYTO9, yellow) and membrane-compromised bacteria and plant nuclei (propidium iodide, magenta). (**e**) Average nodule dry mass and acetylene reduction per nodule dry mass at 24 dpi for plants inoculated with wild-type, Δ*hpnH* or Δ*hpnP*. Data shown in (**b**), (**c**) and (**e**) was collected from n = 8 plants, with error bars representing one standard deviation. Results of two-tailed t-tests between wild type and Δ*hpnH* or Δ*hpnP* are denoted as follows: n.s., p>0.01; **\***, p<0.01; **\*\***, p<0.001; **\*\*\***, p<0.0001.

It is thought that hopanoids primarily promote bacterial survival by rigidifying and decreasing the permeability of membranes (Saenz et al. 2015;, Wu et al. 2015), providing a better barrier against external stress. Structurally distinct hopanoids have different capacities to alter the biophysical properties of membranes and can also differ in the degrees of stress resistance they confer (reviewed in Belin et al. 2018). In the *Bradyrhizobia* genus of legume symbionts, hopanoids promote growth of free-living cultures under acid, salt, detergent, antibiotic, and redox stresses (Silipo et al. 2014; Kulkarni et al. 2015), and we previously showed that these stress resistance phenotypes are largely mediated by the extended hopanoid class (Kulkarni et al. 2015).

We also analyzed an extended hopanoid-deficient mutant of *Bradyrhizobium diazoefficiens* USDA110 in symbiosis with two legumes: the native soybean host for this strain and *Aeschynomene afraspera,* the native host of the closely related photosynthetic *Bradyrhizobia*. *A. afraspera* is a flood-tolerant legume from tropical West Africa, where it has been used in rice intercropping systems (Somado et al. 2003) and to accelerate wound healing in traditional medicine (Swapna et al. 2011; Chifundera 2001; Caamal-Fuentes et al. 2015; Lei et al. 2018). We found that extended hopanoid-deficient mutants of *B. diazoefficiens* fixed less nitrogen per nodule in *A. afraspera* than wild type, while this strain did not appear to have a defect in its native soybean host. Microscopy analyses of a small sample of extended hopanoid mutant-infected *A. afraspera* nodules revealed several aberrant cytological phenotypes, including both nodules containing necrotic signatures, disorganized infection zones, and visible starch granule accumulation (Kulkarni et al. 2015).

These phenotypes are common signatures of poor symbiont performance, yet the lack of genetic tools for *A. afraspera*, the limited literature on this host’s response to non-cooperators compared to model plants, and low number of nodules examined made it difficult to determine the underlying cause. While it has been proposed that hopanoids may enable high rates of symbiotic nitrogen fixation in some hosts by limiting oxygen diffusion across cell membranes (Vilcheze et al. 1994; Parsons et al. 1987; Abeysekera et al. 1990), from our previous assays, we could not determine whether the poor symbiotic performance of extended hopanoid mutants reflects ineffective nitrogen fixation *per se*, or is simply a consequence of lower general stress resistance. Because we did not observe an extended hopanoid mutant phenotype in soybean, we instead suggested that the extended hopanoid mutant may not survive exposure to <u>n</u>odule <u>c</u>ysteine-rich (NCR) peptides, which are synthesized by *A. afraspera* (Czernic et al. 2015) but absent in soybean.

Here, we sought to dissect further the symbiotic phenotypes of *B. diazoefficiens* extended hopanoid mutants in association with *A. afraspera*. We found that the lower nitrogen fixation of extended hopanoid mutants can be fully explained by a reduction in root nodule sizes and rhizobial occupancy, indicating that the underlying defect is unrelated to per-bacteroid nitrogen fixation levels. Using a novel single-nodule tracking approach to quantify *A. afraspera* nodule development, we uncovered both the baseline growth parameters for wild-type nodules and a surprising heterogeneity of extended hopanoid mutant developmental phenotypes. These results challenge the conclusions of our prior study (Kulkarni et al. 2015) and identify new, potentially hopanoid-dependent stages in the *B. diazoefficiens*-*A. afraspera* symbiosis for future mechanistic work. This work also provides a quantitative reference point for understanding the impact of symbiotically ineffective strains on *A. afraspera* nodule development.

## Results

### Loss of extended hopanoids results in reduced nodule size

Previously, we observed a symbiotic defect for an extended hopanoid-deficient (Δ*hpnH*) strain of *B. diazoefficiens* in association with *A. afraspera* (Kulkarni et al. 2015). To further validate this defect, we inoculated *A. afraspera* plants with Δ*hpnH* (lacking extended hopanoids), Δ*hpnP* (lacking 2-Me hopanoids), or wild-type *B. diazoefficiens*. At 24 days post-inoculation (dpi), plants inoculated with Δ*hpnH* were shorter than wild type-inoculated plants, although both strains produced equivalent numbers of nodules (Fig. 1b). Δ*hpnH*-inoculated plants also exhibited a roughly 50% decrease in the rate of acetylene gas reduction compared to wild type-inoculated plants at this time point (Fig. 1c). In contrast, the Δ*hpnP* mutant was similar to wild type (Fig. 1b-c). These results are consistent with our previous findings (Kulkarni et al. 2015).

To assess Δ*hpnH* viability within *A. afraspera* nodules, we performed morphological analyses of nodules using confocal fluorescent microscopy. Fifty-seven wild-type and 67 Δ*hpnH* nodule cross-sections were stained with a bacterial Live:Dead kit, consisting of the cell-permeable SYTO9 dye (staining all cells) and propidium iodide (PI) (staining only cells with a compromised membrane). We did not observe an increase in predominantly PI-stained nodules for Δ*hpnH* compared to wild type (Fig. 1d; Fig. S1, S2). Signatures of plant necrosis, which we previously associated with Δ*hpnH* when we observed a smaller number of nodules (Kulkarni et al. 2015), occurred prominently in only 1/67 Δ*hpnH* nodules examined (Fig. S2).

The most apparent phenotype of Δ*hpnH* nodules was their relatively small size (Fig. S1, S2). We repeated acetylene reduction assays for wild type- and Δ*hpnH*-inoculated plants and then calculated the total nodule dry mass for each plant at 24 dpi. We found a decrease in the nodule dry mass per plant for Δ*hpnH*-inoculated plants that is sufficient to explain the decrease in acetylene reduction rates (Fig. 1e). This result rules out the possibility that nitrogenase functions ineffectively in the absence of extended hopanoids due to inactivation by oxygen, as has been suggested in *Frankia* (Vilcheze et al. 1994; Parsons et al. 1987; Abeysekera et al. 1990), as the per-mg nitrogen fixation rates are not affected by extended hopanoid loss.

### *Δ*hpnH nodules are more variable in size than wild-type nodules

We next measured acetylene reduction per plant across an extended 40 dpi period, and we observed that the differences in both acetylene reduction rates and nodule dry masses between wild type and Δ*hpnH* steadily decreased with time (Fig. 2a-b). By 40 dpi the overall symbiotic efficiencies of wild type and Δ*hpnH* per plant were indistinguishable, in terms of the plants’ qualitative appearance (Fig. 2c-d) as well as their average shoot heights and acetylene reduction rates (Fig. S3). Total nodule counts per plant also did not differ between wild type and Δ*hpnH* at 40 dpi, indicating that the increase in total nodule mass reflects growing nodules rather than more frequent nodulation (Fig. S3).

**Figure 2.**
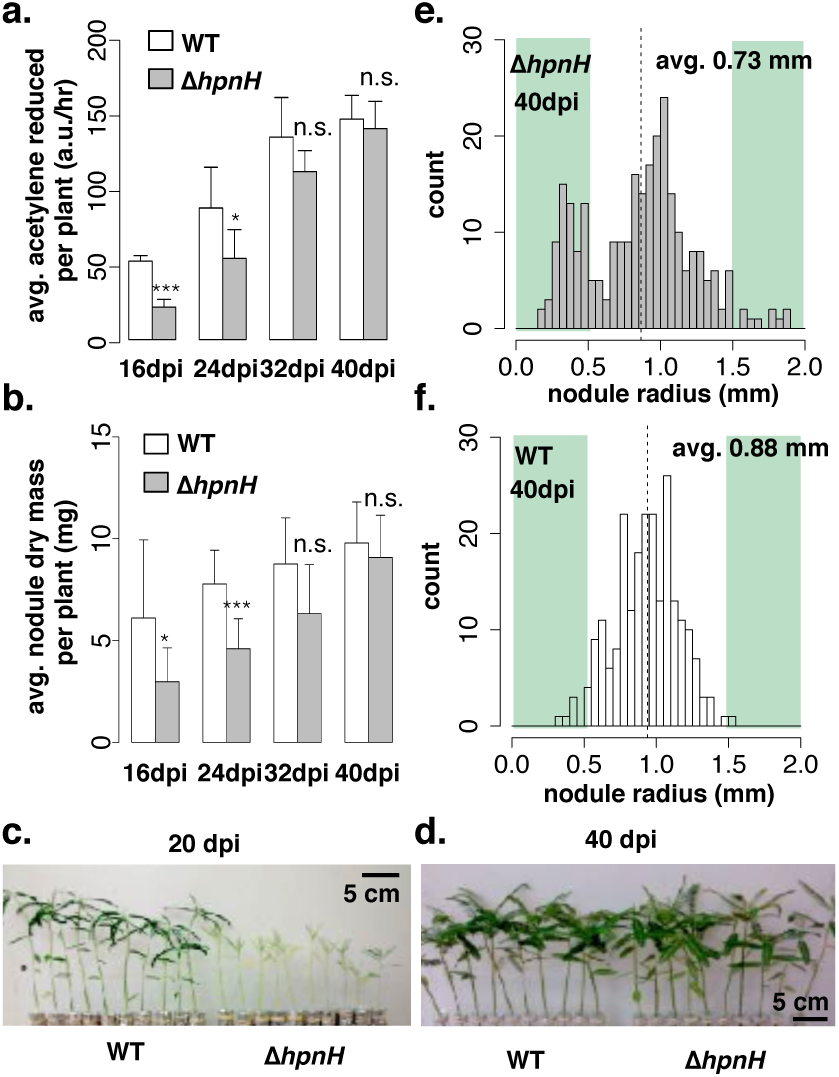
Smaller Δ*hpnH* nodules are offset by increased nodule size heterogeneity over time. (**a**) Average acetylene reduction per plant (n=4 plants per bar) and (**b**) average nodule dry mass per plant (n=8 plants per bar) for *A. asfrapera* inoculated with wild-type or Δ*hpnH* over time. Error bars representing one standard deviation. Results of two-tailed t-tests between wild type and Δ*hpnH* are denoted as follows: n.s., p>0.05; **\***, p<0.05; **\*\*\***, p<0.0001. (**c-d**) *A. afraspera* inoculated with wild type or Δ*hpnH* at (**c**) 20 dpi (left) and at (**d**) 40 dpi (right). (**e-f**) Distributions of nodule diameters at 40 dpi for *A. afraspera* inoculated with (**e**) Δ*hpnH* (right; n=268 nodules pooled from 10 plants) or (**f**) wild type (left; n=227 nodules pooled from 10 plants).

We also measured the radii of individual nodules on ten plants for each strain at 40 dpi (Fig. 2e-f). Interestingly, although *average* nodule sizes did become similar between strains by this time point (0.73 vs. 0.88 mm average radii), their underlying distributions were markedly distinct. Wild-type nodule radii appear to form a roughly normal distribution, whereas the Δ*hpnH* nodule radius distribution is bimodal, consisting of a subpopulation of small nodules with small radii (<0.5 mm) that are rarely observed in wild type, as well as a second, larger subpopulation that has a similar median radius as wild type but is skewed towards larger radii (>1.5mm). These data demonstrate that the small-nodule phenotype of Δ*hpnH* persists throughout a 40 dpi time course, but is compensated by greater size heterogeneity, in which a handful of “mega” nodules offset smaller nodules over time.

### *Δ*hpnH nodule size heterogeneity reflects variable nodule growth rates

To better evaluate the possible origins of the Δ*hpnH* nodule size defect, we studied the kinetics of single nodule development. Beginning one week after inoculation, we collected images of entire plant roots every 3-5 days up to ∼40 days post-inoculation (Fig. S4,S5). From these images, we identified nodules that were clearly visible (*e.g.* not obscured by lateral roots or more recently emerged nodules) in at least five time points (Fig. 3a) and measured their radii. We then calculated nodule volumes by approximating nodules as spheres and plotted the volume of the tracked nodules over time. While we again observed that many Δ*hpnH* nodules were smaller at 40 dpi than any of the wild type nodules, we also found that nodule growth was highly variable both within and between strains (Fig. 3b-c).

**Figure 3.**
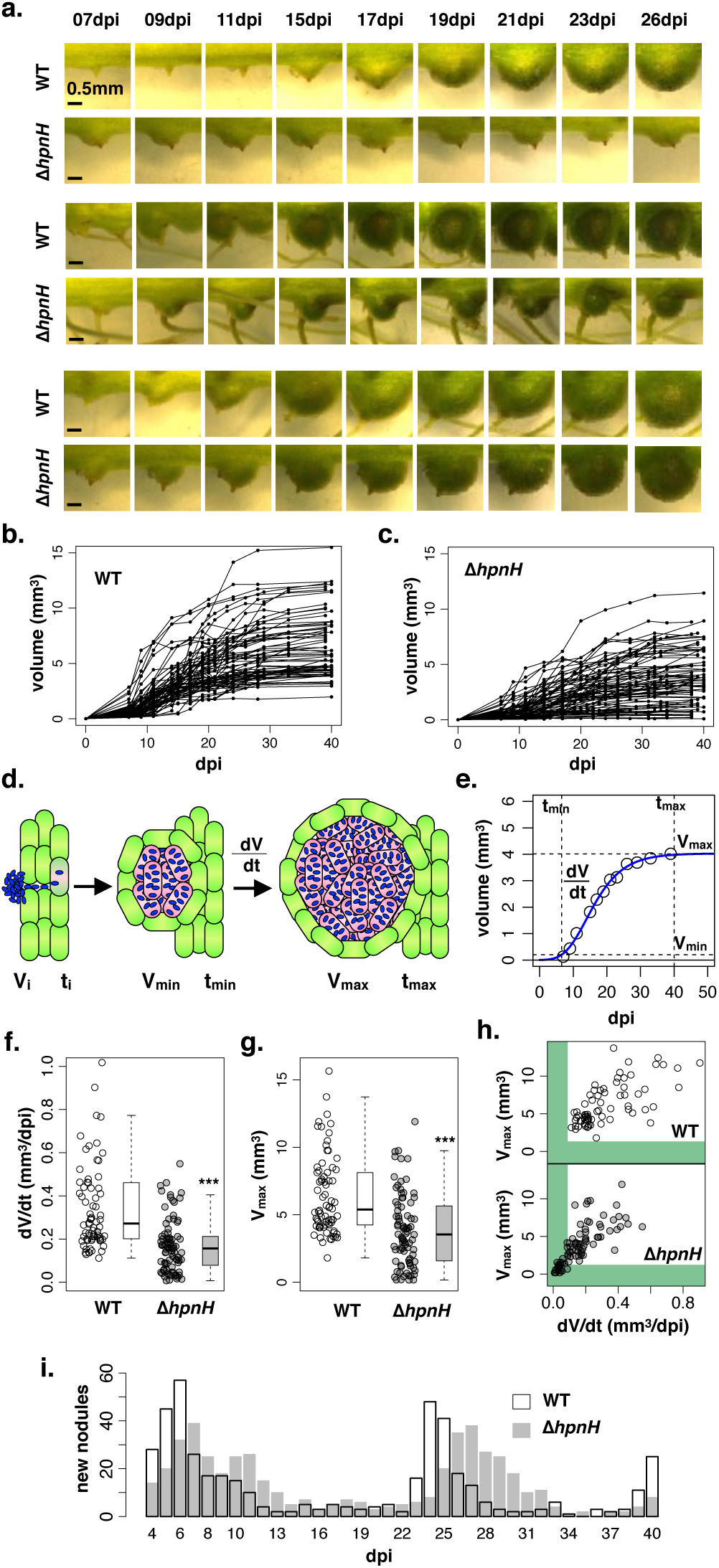
Nodules containing Δ*hpnH* emerge later and have more heterogeneous growth rates and final volumes than wild type. (**a**) Comparison of the development of selected wild type- and Δ*hpnH*-infected nodules over time. (**b**) Nodule growth plots for 74 wild type-infected nodules tracked from 10 plants. (**c**) Nodule growth plots for 84 Δ*hpnH*-infected nodules tracked from 16 plants. (**d**) Schematic of nodule development in *A. afraspera*. From the left, bacteria (in blue) colonize and invade plant roots (green) and intracellularly infect a root cell (pink); the time of this initial intracellular infection is considered **t_i_** and the nodule volume can be described as the volume of the single infected root cell, **V_i_**. This infected cell proliferates to form a spherical nodule that is visible to the naked eye, at time **t_min_** and volume **V_min_**. The infected plant cells continue to proliferate at rate **dV/dt** until the nodule has fully matured at time **t_max_** and volume **V_max_**. (**e**) Fitted growth curve for a sample wild-type nodule illustrating the positions of **t_min_**, **V_min_**, **dV/dt**, **t_max_**, and **V_max_**. (**f-g**) Jitter and box plots of (**f**) **dV/dt** and (**g**) **V_max_** values for all wild type- and Δ*hpnH*-infected nodules. Results of KS-tests between wild-type and Δ*hpnH* nodules are denoted as follows: ***, p<10^−6^. (**h**) Scatter plots of **dV/dt** vs. **V_max_** values for wild-type and Δ*hpnH* nodules. Values of **dV/dt** and **V_max_** below what is observed in the wild-type dataset are highlighted in green. (**i**) Distributions of **t_min_** values (as observed by eye) for nodules from wild type- (white bars) or Δ*hpnH*- (grey bars) infected plants. N=457 wild-type nodules across 20 plants and 479 Δ*hpnH* nodules across 20 plants.

We then developed a simple framework for quantifying nodule development, in which nodule growth is defined by the following variables: the time (*t*_i_) of the initial intracellular infection event and the volume of the nascent nodule (V_i_), equivalent to the volume of one infected *A. afraspera* cortical cell; the time (*t*_min_) and volume (V_min_) at which a clearly visible, spherical nodule has developed; the rate of growth of a nodule once it has become visible (dV/d*t*); and the time (*t*_max_) and volume (V_max_) of a nodule when its growth has stopped (Fig. 3d). To calculate these variables, we fit each nodule’s growth over time to three different growth models: exponential, quadratic, and a generalized logistic (*e.g.* sigmoidal) equation commonly used for plant growth (Szparaga and Kocira 2018; Richards 1959) (see Methods for complete details). Sigmoidal models generally provided the best fit to the experimental data, so these models were used for growth parameter calculation (Fig. 3e; Fig. S6, S7).

The growth rates of Δ*hpnH* nodules were lower on average than wild-type nodules (Fig. 3f), with roughly a third of tracked nodules exhibiting growth rates lower than observed for wild type (<0.1 mm^3^/dpi). A similar fraction of nodules had smaller final volumes than wild type (Fig. 3g). We further found that the growth rate of a nodule and its maximum size are positively linearly correlated for both strains, with Pearson coefficients of ∼0.64 (p<10^−9^) for wild type and ∼0.75 (p<10^−15^) for Δ*hpnH,* and that the subpopulation of nodules with lower-than-wild-type growth rates and small nodule sizes are the same (Fig. 3h). We interpret these data to suggest that host proliferation is slower in a subset of nodules infected with Δ*hpnH*, and that this largely accounts for the low final volume of these nodules.

We also noted that Δ*hpnH* nodule sizes at 40 dpi differed between these single-nodule volume measurements (Fig. 3g) and our previous 40 dpi end-point measurements of nodule radii (Fig. 2e), in that we did not observe larger-than-wild-type “mega” nodules in the single-nodule dataset. This discrepancy likely reflects the smaller sample size in our single-nodule tracking experiments (84 compared to 268 end-point nodules), and the low frequency of “mega” nodule formation. To verify this, we selected 10,000 random subsets of 84 nodules from the 268 Δ*hpnH* nodules shown in Figure 2e, converted the nodule radii to volumes, and found that there is no statistically significant difference (p<0.05) between a random subset of Fig. 2e and the Δ*hpnH* single-nodule tracking data in ∼92% (9184/10000) of cases. Thus the differences in nodule size distributions in Figure 3g and Figure 2e are consistent with sampling error.

We also calculated each nodule’s window of maximum growth, defined as the time required for a nodule to increase from 10% to 90% of its final volume. Neither the time at which a nodule reaches 90% of its maximum volume, t_max_, nor the window of maximum growth differs significantly between Δ*hpnH* and wild type (Fig. S8a-b). The window of maximum growth for each nodule is also uncorrelated with their final volume or growth rate, indicating that small nodules are not prematurely aborted; rather, their growth periods are similar to larger nodules (Fig. S9a-d).

To better understand the subpopulation of small, slow-growing Δ*hpnH* nodules, we isolated nodules with <0.5 mm radius, sectioned and stained them with SYTO9, PI and Calcofluor, and imaged them with confocal microscopy. We found that while most small Δ*hpnH* nodules contained a single, continuous infection zone, a large fraction were un- or under-infected with bacteria, often exhibiting disorganized central infection zones (∼37%; 28/75) (Fig. 4a; Fig. S10). Of the fully infected small Δ*hpnH* nodules, a subset contained primarily PI-stained, likely dead bacterial cells (∼25%; 12/47) (Fig. 4a; Fig. S10). Similar proportions of under-infected nodules or nodules primarily occupied with membrane-compromised bacteria did not occur in larger Δ*hpnH* nodules harvested at the same time point, although fragmented infection zones were still common (Fig. 4b; Fig. S11). We also compared the subpopulation of small Δ*hpnH* nodules at 40 dpi to two wild-type nodule populations: similarly small nodules harvested at 10 and 25 dpi (Fig. 4a; Fig S12; Fig. S13), and nodules harvested at the same 40 dpi time point (Fig. 4b; Fig. S14). Again, we found that high proportions of under-infected nodules and membrane-compromised bacteria were unique to the Δ*hpnH* small-nodule subset.

**Figure 4.**
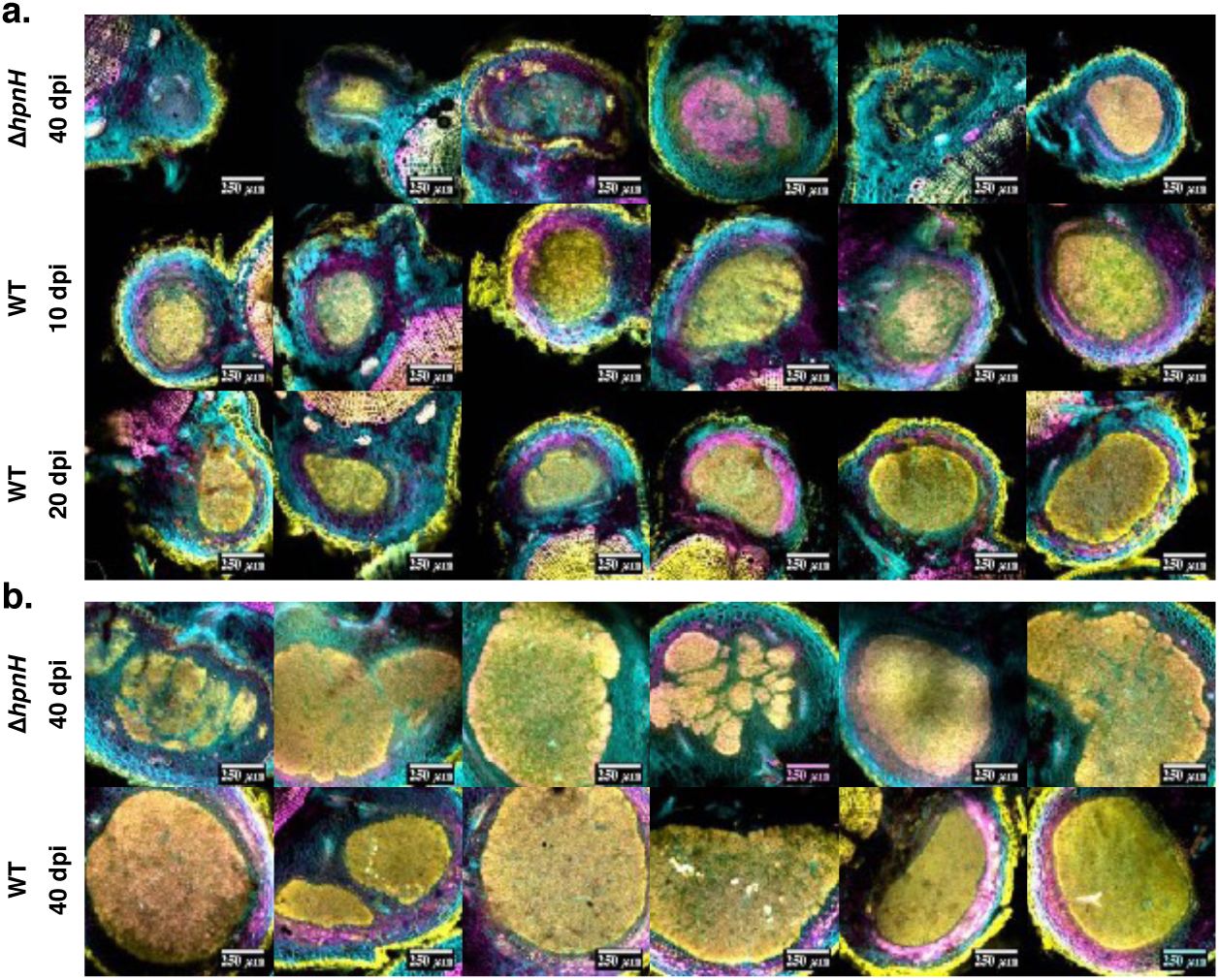
Small Δ*hpnH* nodules are under-infected compared to wild type. (**a**) Confocal sections of small (<0.5 mm radius) Δ*hpnH*-infected nodules harvested at 40 dpi and small (<0.5 mm radius) wild type-infected nodules harvested at 10 and 25 dpi. (**b**) Confocal sections of larger (>0.5 mm radius) Δ*hpnH*- or wild type-infected nodules harvested at 40 dpi.

### *Δ*hpnH nodule emergence is delayed

The “true” beginning of nodule formation is the time when the first *A. afraspera* cortical cell is infected, *t*_i_ (Fig. 3d). However, this initial infection event is not visible at the root surface, and it is difficult to extrapolate from sigmoidal models in which the growth curves approach the initial volume V_i_ ∼ 0 mm^3^ asymptotically. As a proxy for *t*_i_, we defined three alternate t_min_ as the times at which nodules reached three arbitrarily small volumes: V = 0.05 mm^3^, V = 0.1 mm^3^, and V = 0.2 mm^3^. When t_min_ is defined by V = 0.05 mm^3^ or 0.1 mm^3^, t_min_ could not be accurately calculated for all nodules, as the sigmoidal models sometimes predicted an impossible t_min_ < 0 (Fig. S8c). These nodule volumes are also too small to be seen on the root surface, and we had no experimental means to determine the accuracy of the calculations in this low-volume regime. When t_min_ is defined by V = 0.2 mm^3^ (the smallest nodule volume that we could identify in our single-nodule tracking assays), there is a small but statistically significant increase for Δ*hpnH* relative to wild type (Fig. S8c).

To independently verify this delay in nodule emergence, we inspected the roots of 20 wild type- and 20 Δ*hpnH*-inoculated plants over 40 dpi and recorded the number of visible nodules per plant each day. We found a more even distribution of observed t_min_ for Δ*hpnH* relative to wild type, with a 1-3 day shift in the most frequent dpi. Surprisingly, we also found that the formation of new nodules is periodic, with a new “burst” of nodules emerging roughly every 18 days (Fig. 3i). This periodicity of nodule emergence appears to be similar between strains.

While the slight t_min_ delay for Δ*hpnH* is consistent with longer times required to initiate the symbiosis (*e.g.* root surface colonization, invasion of the root epidermis and cortex, and intracellular uptake), it is also possible that a delay in t_min_ simply reflects a lower rate of nodule growth immediately after the first intracellular infection. To address this, we compared the calculated value of t_min_ (defined by V = 0.2 mm^3^) to the maximum growth rates and volumes for each nodule (Fig. S9e-f). We did not find that nodules with lower growth rates and final volumes than wild type were more likely to have a later t_min_, supporting the interpretation that the delay in t_min_ of Δ*hpnH* could be due to a separate initiation defect. Interestingly, t_min_ is also not correlated with the period in which maximum nodule growth occurs, such that later-emerging nodules have similar growth period to nodules formed within a few dpi (Fig. S9g-h). This indicates that although nodule emergence is restricted to narrow, periodic windows (Fig. 3i), once a nodule has entered its maximum growth phase, its continued growth is comparatively unconstrained.

### *Δ*hpnH is delayed in a pre-endosymbiont stage

We next performed competition assays using a standard fluorescence labeling approach. We first generated Δ*hpnH* and wild-type strains expressing chromosomally-integrated fluorescent proteins, and then we co-inoculated *A. afraspera* with different ratios of these two strains. As control experiments, we also co-inoculated each tagged strain with its untagged counterpart, in order to determine the effect of fluorescent protein overexpression on each strain’s competitiveness. After 40 dpi we measured the size of nodules on plants inoculated with each strain combination and ratio, then sectioned and fixed nodules for imaging. Although we expected each nodule to contain a clonal population of symbionts based on previous work (Bonaldi et al. 2011; Ledermann et al. 2015), the majority of nodules instead contained a mixture of both strains (Fig. 5a).

**Figure 5.**
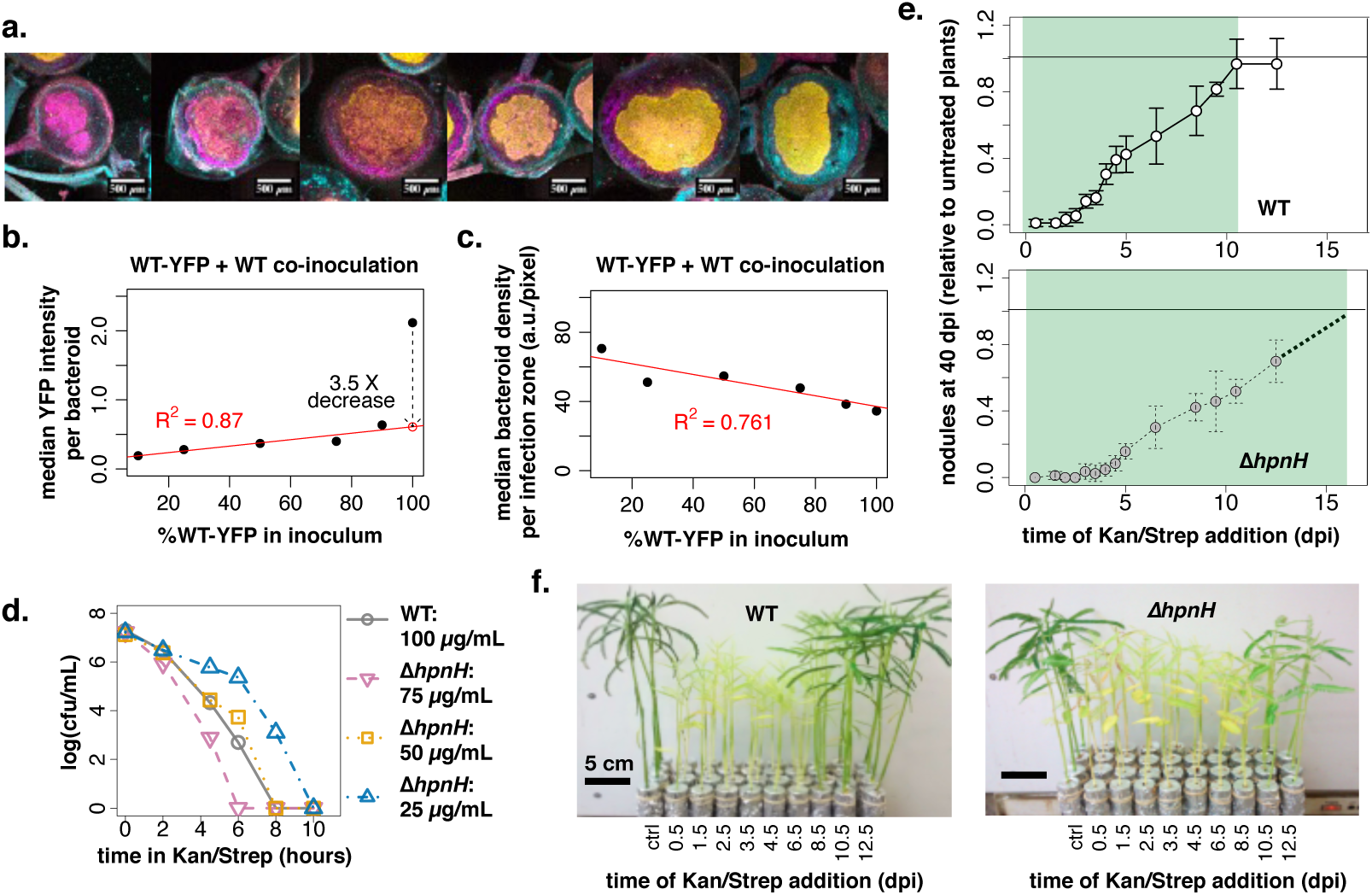
Extended hopanoid mutants are delayed at pre-intracellular stage(s) in symbiosis development. (**a**) Confocal sections of nodules from plants co-inoculated with wild type-YFP and Δ*hpnH*-mCherry harvested at 45-55 dpi. Sections were stained with Calcofluor (cyan) and are expressing YFP (yellow) and mCherry (magenta). (**b**) Scatter plot of median YFP intensity per pixel normalized by propidium iodide intensity per pixel (*e.g.* bacteroid density) within infection zones of nodules from plants co-inoculated with wild type-YFP and wild type, as a function of the percentage of wild type-YFP in the inoculum. (**c**) Scatter plot of median propidium iodide intensity per pixel (*e.g.* bacteroid density) within infection zones of nodules from plants co-inoculated with YFP-tagged wild type and untagged wild type, as a function of the percentage of WT-YFP in the inoculum. (**d**) Colony forming units/mL in wild type and Δ*hpnH* cultures grown in BNM supplemented with varying concentrations of kanamycin and spectinomycin at various times post-inoculation. (**e**) Average nodules per plant at 40 dpi for plants inoculated with either wild type or Δ*hpnH* and treated with 50 µg/mL (Δ*hpnH*) or 100 µg/mL (wild type) kanamycin and streptomycin at various time points post-inoculation. Nodule counts are normalized to those observed in non-antibiotic treated plants. (**f**) Images of inoculated plants at 40 dpi after antibiotic treatment at various time points. Untreated plants are shown on the left, with increasing time of antibiotic addition. Error bars represent one standard deviation.

We quantified the relative abundance of each strain in each nodule by fluorescence imaging; in our control experiments, in which only one fluorophore-expressing strain was present, a DNA dye was used to label all bacteria. Both WT-YFP and Δ*hpnH*-mCherry were significantly out-competed by their corresponding untagged strains, with higher proportions of tagged strains correlating with lower bacterial DNA abundance and smaller nodule and/or infection zone sizes (Fig. 5b-c; Fig. S16-S18). Additionally, plants co-inoculated with untagged-Δ*hpnH* and Δ*hpnH*-mCherry were significantly shorter than plants inoculated with untagged-Δ*hpnH* only, suggesting Δ*hpnH*-mCherry is symbiotically defective (Fig. S15).

These effects of fluorophore overexpression made it difficult to interpret our WT-YFP and Δ*hpnH*-mCherry competition data, so we developed an alternative, antibiotics-based method to study the timing of early symbiotic initiation. First, we identified antibiotics that were effective against *B. diazoefficiens* but would minimally affect *A. afraspera* growth. We tested three antibiotics (100 µg/ml streptomycin, 100 µg/ml kanamycin, and 20 µg/ml tetracycline) and treated non-inoculated plants with these antibiotics for two weeks, alone and in combination. After this treatment, we found that neither kanamycin nor streptomycin, nor the combination of the two, significantly affected plant appearance, shoot height, or root and shoot dry masses compared to untreated controls (Fig. S19). Plants treated with tetracycline were noticeably more yellow in color, indicating chlorosis, and the roots and plant medium became brown; these plants also had lower shoot and root dry masses than untreated plants (Fig. S19).

Next, because the Δ*hpnH* strain is more sensitive to antibiotics than wild type (Kulkarni et al. 2015), we tested various concentrations of the non-plant-perturbing antibiotics streptomycin and kanamycin to identify concentrations that would result in the same rates of cell death for both strains. We inoculated plant growth media with wild type or Δ*hpnH* to the same cell densities and under the same environmental conditions as in plant inoculation experiments. The wild-type culture was supplemented with 100 µg/ml streptomycin plus 100 µg/ml kanamycin, and Δ*hpnH* cultures were supplemented with decreasing concentrations of these antibiotics: 75, 50 and 25 µg/mL each. Samples of the cultures were then collected, serially diluted and added to PSY plates to estimate colony-forming units (cfus) per mL over time. At 50 µg/mL kanamycin plus 50 µg/mL streptomycin, the rate of decrease in cfus/mL for Δ*hpnH* was equivalent to that of wild type treated with 100 µg/ml kanamycin plus streptomycin (Fig. 5d).

Finally, we inoculated 40 plants each with wild type or Δ*hpnH* and added streptomycin or kanamycin to 100 µg/mL each or 50 µg/mL each, respectively, at various points post-inoculation. After 40 days we counted the number of nodules per plant, and found that antibiotics were able to block nodule formation over a ∼50% longer window in Δ*hpnH* compared to wild type (Fig. 5e). The decrease in nodules formed at different antibiotic treatment time points was also evident in the overall appearance of the plants (Fig. 5f). These results suggest that Δ*hpnH* requires more time on average to reach the intracellular stage of the symbiosis, at which point we presume that the bacteria are protected from antibiotic by the host cells. These data are consistent with Δ*hpnH* requiring more time to colonize the root surface, invade the root epidermis, and/or be internalized by host cells.

### Extended hopanoids support surface attachment and motility in vitro

Because we found that expression of genetic tags in wild type and Δ*hpnH* perturbed their symbiosis with *A. afraspera,* and because we found that the hopanoid mutant viability is reduced by sonication, centrifugation, and mechanical or detergent-based tissue disruption techniques required to re-isolate bacteria from plants, we could not confidently follow these strains *in planta*. Instead, we used an *in vitro* approach to study two steps in the initiation of the symbiosis: (1) bacterial motility toward or along the *A. afraspera* root, and (2) stable attachment of bacteria to the root surface (Wheatley and Poole 2018). To determine whether Δ*hpnH* is less motile than wild type, we inoculated low-agar, PSY plates with Δ*hpnH* or wild type and measured the rate of zone of swimming over time. We observed that diameter of motility was reduced in Δ*hpnH* compared to wild type (Fig. 6a-b), consistent with a swimming motility defect; however, because we have previously shown that Δ*hpnH* grows more slowly in this medium than wild type (Kulkarni et al. 2015), we could not rule out the possibility that slower zone expansion simply reflects a longer doubling time.

**Figure 6.**
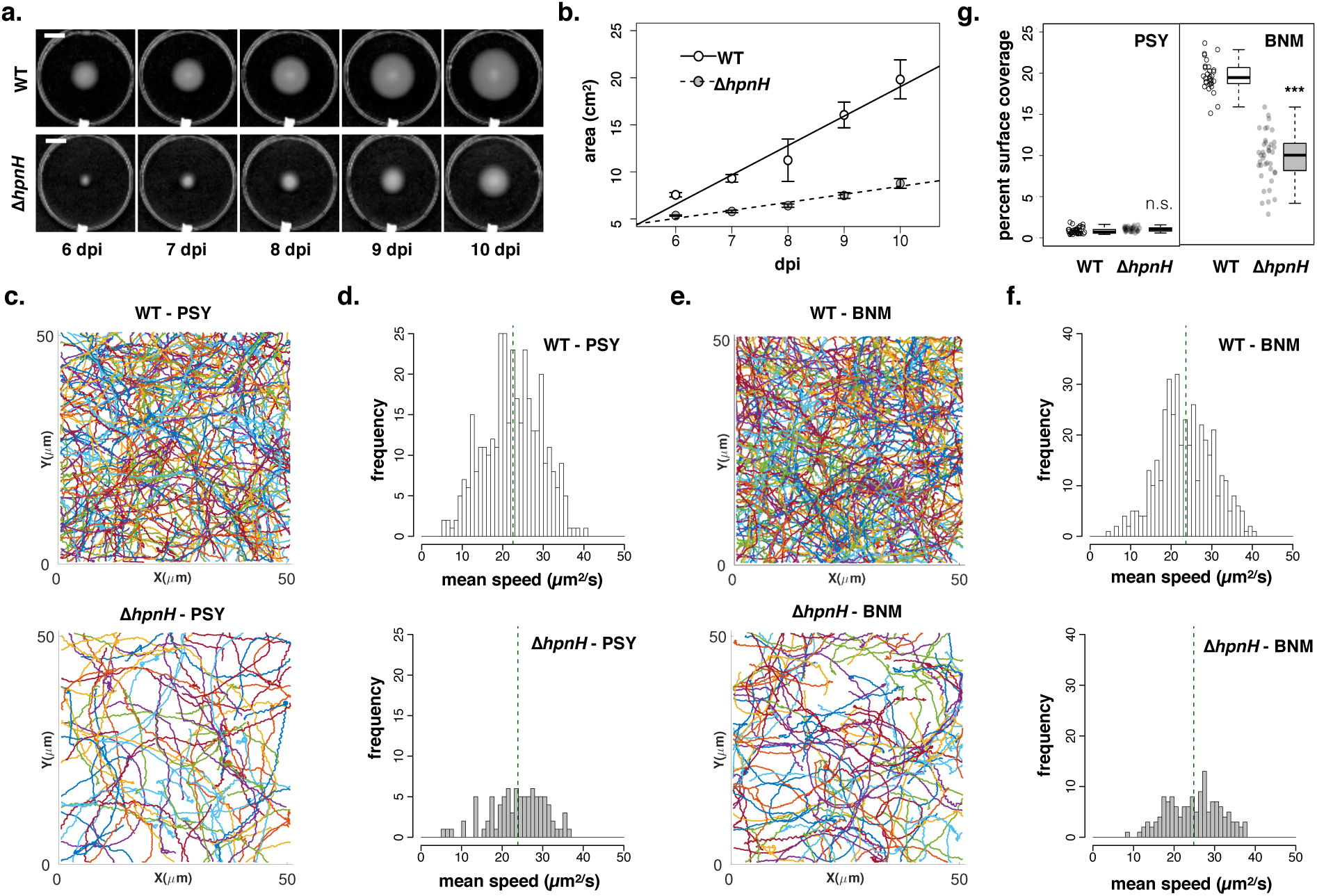
Extended hopanoid mutants are less motile than wild type and attach poorly to surfaces *in vitro*. (**a**) Sample time course of wild type and Δ*hpnH* colony expansion on low-agar PSY plates (dpi = days post-inoculation). Scale bars represent 2 cm. (**b**) Average colony sizes of wild type and Δ*hpnH* over time. N=4 plates per strain; error bars indicate one standard deviation. (**c**) Trajectories of individual wild type (top) and Δ*hpnH* (bottom) cells over a 5 minute time course in PSY. (**d**) Distributions of mean-speed s for motile wild type (N=359) and Δ*hpnH* (N=91) cells for trajectories in **d**. Dotted lines indicate the means of the distributions. (**e**) Trajectories of individual wild type (top) and Δ*hpnH* (bottom) cells over a 5 minute time course in BNM. (**f**) Distributions of mean-speeds for motile WT (N=421) and Δ*hpnH* (N=141) cells in BNM for trajectories in **e**. Dotted lines indicate the means of the distributions. (g) Jitter and box plots of surface attachment (*e.g.* the percent of the field of view covered with cells) of WT and Δ*hpnH* after 2 hours of incubation on glass in PSY or BNM. N=40 fields of view per condition. Results of two-tailed t-tests between wild type and Δ*hpnH* are denoted as follows: n.s., p>0.05; **\*\*\***, p<0.00001.

To investigate the nature of the plate motility defect, we studied the motility of single *B. diazoefficiens* cells. We inoculated cells into a glass-bottom, sterile PSY flow cells with 100µL of each strain and recorded the movement of cells near the glass surface at 5 ms time resolution. Trajectories of individual swimming cells, defined as having super-diffusive motion and a trajectory radius of gyration >2.5 µm, were calculated and analyzed in MATLAB (Lee et al. 2018). In agreement with our low-agar swimming plates, we found significantly fewer (p < 0.0001) motile cells for Δ*hpnH* (N = 65 ± 29) than wild type (N= 368 ± 60) in PSY medium (Fig. 6c; Table S1). The average mean-speed among motile cells were similar between strains: <V>_ΔhpnH_ = 24.83 ± 7.0 µm/sec and <V>_wt_ = 22.75 ± 6.7 µm/sec (Fig. 6d; Table S1). Because the composition of PSY differs greatly from that of the plant growth medium (BNM), we repeated these assays in BNM supplemented with arabinose. Under this condition, we again observed a lower fraction of motile Δ*hpnH* cells than wild type (N_ΔhpnH_ = 54 ± 59, N_wt_= 450 ± 310) with similar mean speeds between strains (Fig. 6e-f; Table S1).

We next tested the surface attachment capabilities of Δ*hpnH* and wild type by incubating dense bacterial cultures on glass coverslips and quantifying the fraction of the surface covered with stably adherent cells after two hours. In PSY medium, both strains adhered poorly, and there was no significant difference in their attachment efficiencies (Fig 6g; Fig. S20). In BNM supplemented with arabinose, both strains adhered to glass better than in PSY, and Δ*hpnH* attachment levels were significantly lower than wild type (Fig. 6g; Fig. S20). The decreased adhesion and reduced motile cell population of Δ*hpnH* suggest that stable root colonization by this strain may be less efficient.

## Discussion

Hopanoids are well-established mediators of bacterial survival under stress, and previously we showed that the capacity for hopanoid production is enriched in plant-associated environments (Ricci et al. 2014) and required for optimal *Bradyrhizobia*-*Aeschynomene* spp. symbioses (Silipo et al. 2014; Kulkarni et al. 2015). Here we performed a detailed, quantitative evaluation of the extended hopanoid phenotypes in the *Bradyrhizobium diazoefficiens*-*Aeschynomene afraspera* symbiosis. We determined that extended hopanoid mutants fix nitrogen at similar rates as wild type on a per-bacteroid level, demonstrating that in this host, extended hopanoids are not required to protect nitrogenase from oxygen, as often has been speculated (reviewed in Belin et al. 2018). Instead, we found that the extended hopanoid mutants’ lower *in planta* productivity can be fully attributed to changes in the kinetics of nodule development. By tracking the development of individual root nodules, we observed later nodule emergence times in Δ*hpnH*-inoculated plants. *In vitro*, Δ*hpnH* cells adhered poorly to glass and were less motile than wild type, suggesting they may colonize roots less efficiently (Fig. 7a-b). A third of Δ*hpnH* nodules also grew significantly slower than wild type and were smaller at maturity. Many of these small nodules contained low symbiont densities; a subset of larger Δ*hpnH* nodules also had lower symbiont loads, due to infection zone fragmentation. The origin of this under-infection is unclear. It is possible that bacteria are inefficiently internalized or retained, and this phenotype is simply propagated as nodules develop (Fig. 7c-d). Alternatively, low symbiont densities may reflect symbiont degradation in a previously fully infected nodule (Fig. 7e).

**Figure 7.**
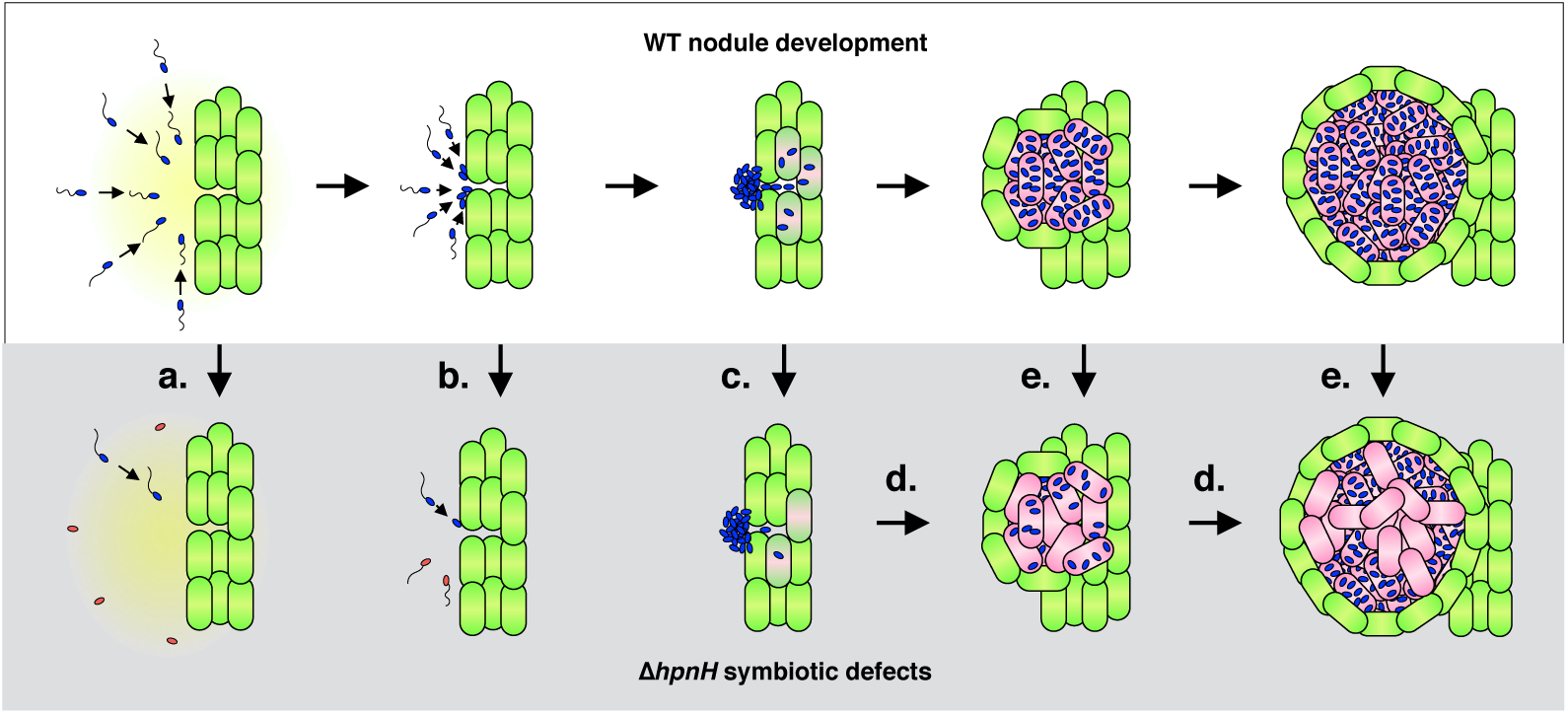
Consequences of extended hopanoid loss in *A. afraspera* nodule development. Schematic representation of *A. afraspera* wild-type root nodule development (top row; white background) and defects in development associated with extended hopanoid loss (bottom row; grey background). Early in development, fewer Δ*hpnH* cells are motile (**a**) and competent to attach to root surfaces (**b**), leading to a delay in establishment of stable root colonies. At later stages, slow growth of Δ*hpnH* into the root interior, or poor uptake by and division within host cells (**c**) may generate “patchy”, or under-populated infection zone that is propagated as the nodule grows (**d**). Alternately, fully-infected Δ*hpnH* nodules may lose symbionts to symbiont cell death (**e**) via poor bacteroid survival or plant-directed symbiosome degradation.

These observations challenge two conclusions from our previous work, requiring a refinement of our interpretation of the roles of extended hopanoids in the plant context (Kulkarni et al. 2015). First, we reported that there was no symbiotic defect of the Δ*hpnH* strain in soybean, based on the observation that nitrogen fixation per mg nodule dry weight was similar to wild type. Given that this study revealed that a reduction in nodule dry weight explains the Δ*hpnH* defect in *A. afraspera*, it is possible that this strain is also defective in its native host, but this defect was obscured by differences in normalization between the soybean and *A. afraspera* datasets. Second, the majority of Δ*hpnH* nodules had wild type-like growth kinetics and morphologies, with a few “mega” nodules displaying unusually fast growth. This finding appears inconsistent with an inability to survive NCR peptides, unless NCR peptide expression levels are extremely variable, or if the mechanisms that compensate for extended hopanoid loss are inconsistent.

What other mechanisms might underpin these extended hopanoid mutant phenotypes? Perhaps they are simply consequences of less rigid *B. diazoefficiens* membranes. The fraction of motile cells in *E. coli* populations has been suggested to be sensitive to changes to the mechanical properties of the outer membrane (Gupta et al. 2006), and membrane-based mechanotransduction is required by diverse bacteria to stimulate extracellular matrix production and cement their attachment to surfaces (Petrova and Sauer 2012; Persat 2017). *B. diazoefficiens* mutants with weakened cell walls also have been shown to be deficient in symbiosis with *A. afraspera* through an NCR peptide-independent mechanism (Barriere et al. 2017), which may be elicited by Δ*hpnH*. Extended hopanoid loss may also have secondary effects on *Bradyrhizobium*-*Aeschynomene* signaling. In the *Frankia*-actinorhizal symbiosis, bacterial extended hopanoids can contain the auxinomimetic compound phenyl-acetic acid (PAA) (Hammad et al. 2003), and though the effects of hopanoid loss on the bacterial metabolome have not been examined, changes in hopanoid production may impact the synthesis and/or secretion of symbiotically active compounds. Future work will be required to determine whether changes in signaling or membrane mechanics dominate the hopanoid mutant phenotypes, and at which developmental stages.

Regardless of the underlying mechanism, it is curious that the absence of extended hopanoids is not a death knell for the *B. diazoefficiens-A. afraspera* symbiosis at any stage. In our *in vitro* studies, mean speeds among motile Δ*hpnH* cells were indistinguishable from wild type, and though we cannot rule out more subtle defects in the direction of movement or chemotaxis, this suggests that motility systems of Δ*hpnH* cells function properly once induced. Similarly *in planta*, Δ*hpnH* nodules developing at wild-type rates and reaching average wild-type volumes did occur – and, in the case of “mega” nodules, some exceeded their wild-type counterparts.

Why do Δ*hpnH* populations form two distinct populations (wild-type-like or defective) rather than falling on a continuous distribution of behavior? Bimodality can reflect switch-like, or threshold-based, regulation, and perhaps in the Δ*hpnH* strain, a fraction of cells cannot support levels of signaling above the threshold required for proper function. Nodules may also differ in the extent to which extended hopanoid loss is compensated. In *Methylobacterium extorquens* and *Rhodopseudomonas palustris* (Bradley et al. 2017; Neubauer et al. 2015), hopanoid loss results in upregulation of other membrane-rigidifying lipids including carotenoids and cardiolipins, and in other plant-microbe systems, lipid exchange between hosts and microbes has been observed (Keymer 2018), suggesting that Δ*hpnH* nodule phenotypes may relate to the local availability of structurally or functionally similar metabolites. Because of these diverse possible explanations for Δ*hpnH* heterogeneity, a detailed comparison of wild type-like and defective nodules, including the distributions of lipids and other metabolites, bacteroid morphology and penetrance, and gene expression variability, will be required to determine why some Δ*hpnH* nodules succeed and others do not.

Beyond hopanoids, our results provide insight into the developmental control of nodule formation by *A. afraspera* hosts. We find that nodulation occurs in bursts separated by fixed 18-day intervals, and that the timing of these bursts is unrelated to net fixed nitrogen production across the root, more likely reflecting the inherent dynamics of the underlying signaling networks. The growth period of individual nodules is similarly deterministic, suggesting that *A. afraspera* hosts do not respond to ineffective symbionts by prematurely aborting nodule development. Rather, we find that *A. afraspera* nodules can be primarily distinguished by their growth rates, *e.g.* the frequencies of infected host cell division. This finding suggests that in *A. afraspera* host cell mitosis and symbiont performance may be coupled, enabling future studies on the molecular signals through which this coupling occurs.

Finally, our results underscore the importance of identifying the most informative, least perturbing tools for interrogating legume-microbe symbiosis. Employing quantitative, time-resolved, single-nodule and single-cell approaches rather than bulk measurements were essential for uncovering the diverse phenotypes of the *B. diazoefficiens* extended hopanoid mutants and yielded unexpected information on regulation of nodule development by *A. afraspera*. We have also shown the limitations of introducing overexpressed genetic tags into bacteria. While use of these tags has undoubtedly enhanced our understanding of legume-microbe symbiosis (Ledermann et al. 2018), they may not fully capture the behavior of native organisms. Additionally, our work is one of many to emphasize the importance of appropriate culture models for mimicking the host environment, as the Δ*hpnH* surface attachment defect was observed in plant growth medium but not in a standard richer medium. A more detailed analysis of the host environment, including the full milieu of root exudates (Sugiyama and Yazaki 2012), available carbon sources (Pini et al. 2017) and trace metals specific to each legume, will improve *in vitro* models of legume-bacteria interactions and may allow selection of strains with improved performance in agriculture.

## Methods

### B. diazoefficiens culture and strain generation

*B. diazoefficiens* hopanoid biosynthesis mutants were generated previously (Kulkarni et al. 2015). For construction of YFP- and mCherry-expressing strains, fluorophore expression vectors pRJPaph-YFP and pRJPaph-mCherry (Ledermann et al. 2015) were provided as a gift from Prof. Dr. Hans-Martin Fischer (ETH Zurich). These vectors were introduced into *B. diazoefficiens* by conjugation with the β2155 DAP auxotroph strain of *E.coli*, using the following protocol: *B. diazoefficiens* wild type and Δ*hpnH* were grown in 5 mL PSY medium(Regensburger and Hennecke 1983) at 30°C and 250 rpm to an OD_600_ of ∼1.0 (wild type) or of 0.5-0.8 (Δ*hpnH*). β2155 strains carrying pRJPaph vectors were grown to an OD_600_ of 0.5-0.8 in 5 mL LB supplemented with 10 µg/mL tetracycline and 300 µm DAP at 37°C and 250 rpm. Both *B. diazoefficiens* and β2155 donor cultures were pelleted at 3250 x g for 30 minutes, washed three times in 0.9% sterile saline, and resuspended in 0.9% sterile saline to a final OD_600_ of 1.0. *B. diazoefficiens* strains and β2155 donor cells were combined at a 4:1 ratio, respectively, and mixed by repeated pipetting. Aliquots (50 µl) of these 4:1 mixtures were dropped to PSY plates supplemented with 300 µm DAP, dried in a biosafety cabinet, and incubated for 48 hours at 30°C. Conjugation pastes were then removed from plates and resuspended in 5 mL sterile saline, pelleted at 3250 x g for 30 minutes and washed twice, in order to remove residual DAP. Washed cells were pelleted a final time and resuspended to 200 µl in 0.9% sterile saline and plated onto PSY plates supplemented with 20 µg/mL (wild type) or 10 µg/mL (Δ*hpnH*) tetracycline. Colonies appeared after 7-10 days (wild type) or 10-14 days (Δ*hpnH*) and were streaked onto fresh PSY/tetracycline plates, then screened for fluorescence using a Lumascope 720 fluorescent microscope (Etaluma). Fluorescent colonies were then sequenced to verify insertion of the pRJPaph vectors into the *scoI* locus.

### A. afraspera *cultivation and inoculation with* B. diazoefficiens

*A. afraspera* seeds were obtained as a gift from the laboratory of Dr. Eric Giraud (LSTM/Cirad, Montpelier, France). Seeds were sterilized and scarified by incubation in 95% sulfuric acid at RT for 45 minutes, followed by 5 washes in sterile-filtered nanopure water and a second incubation in 95% ethanol for 5 minutes at RT. After ethanol treatment seeds were washed 5X and incubated overnight in sterile-filtered nanopure water. Seeds were then transferred to freshly poured water/agar plates using sterile, single-use forceps in a biosafety cabinet, and germinated for 24-72 hours in the dark at 28-32°C.

Seedlings were then placed in clear glass test tubes containing 100 mL of sterile, nitrogen-free Buffered Nodulation Medium (BNM)(Ehrhardt et al. 1992) and grown for 7-10 days in plant growth chambers (Percival) under the following settings: 28°C, 80-90% humidity, and 16 hour photoperiod under photosynthetic light bulbs (General Electric) emitting ∼4000 lumens/ft^2^. In parallel, *B. diazoefficiens* strains were grown in 5-10 mL PSY liquid culture at 30°C and 250 rpm to stationary phase (OD_600_ > 1.4). Stationary phase cultures were diluted into PSY one day prior to plant inoculation to reach an OD_600_ of ∼0.8 at the time of inoculation. OD_600_ ∼ 0.8 cultures were pelleted at 3250 x g for 30 minutes at RT, washed once in PSY, then resuspended in PSY to a final OD_600_ of 1.0. Resuspended *B. diazoefficiens* cultures were directly inoculated into the plant medium in a sterile biosafety cabinet; 1 mL of OD_600_=1.0 culture was added per plant. Inoculated plants were then returned to growth chambers and maintained for the times indicated for each experiment. For longer experiments (lasting longer than ∼30 days post-inoculation), plant growth tubes were refilled with sterile-filered nanopure water as needed. To minimize cross-contamination, inoculated plants and non-inoculated plants were cultivated in separate growth chambers, and growth chambers were sterilized with 70% ethanol followed by UV irradiation for at least 24 hours between experiments.

### Acetylene reduction experiments

Individual plants were transferred to clear glass 150 mL Balch-type anaerobic culture bottles containing 15 mL BNM medium and sealed under a gas-tight septum. After sealing, 15 mL of headspace gas (10% of the culture bottle volume) was removed and replaced with 15 mL of acetylene gas (Airgas). Plants in culture bottles were incubated in the light at 28°C in growth chambers for 3-6 hours. A 100 µl sample of the headspace gas was removed using a gas-tight syringe (Hamilton), and this sample was injected and analyzed for ethylene signal intensities using a Hewlett Packard 5890 Series II GC with Hewlett Packard 5972 Mass Spectrometer with a 30mx0.320mm GasPro Column (Agilent Technologies) and a 2 mm ID splitless liner (Restek Corporation). Following acetylene reduction measurements, plants were removed from jars and plant shoot heights and number of nodules per plant were recorded. When nodule dry mass measurements were performed, nodules were harvested with a razor blade, transferred into pre-weighed Eppendorf tubes, dried at 50°C for a minimum of 48 hours, then weighed again.

### Live:Dead staining and imaging of nodule cross-sections

Nodules were hand-sectioned with razor blades and immediately transferred into a fresh solution of 5 µM SYTO9 (diluted 1:100 from a 500 uM stock in DMSO at −20°C; Thermo Fisher) and 0.02 mg/mL (30 µM) propidium iodide (diluted 1:50 from a 1 mg/mL stock stored in water at 4°C; Thermo Fisher) in PBS. Nodule sections were incubated in this SYTO9/propidium iodide solution at room temperature for 30 minutes in the dark with gentle shaking, washed 5X in PBS, and fixed in 4% paraformaldehyde (Electron Microscopy Sciences) in PBS overnight in the dark at 4°C. Fixed sections were washed 5X in PBS and transferred to a freshly prepared solution of 0.1 mg/mL Calcofluor White (Fluorescence Brightener 28; Sigma) in PBS. The sections were incubated in the Calcofluor solution in the dark for 1 hour at RT with gentle shaking and washed 5X in PBS to remove excess dye.

Prior to imaging, sections were transferred to 30 mm imaging dishes with 20 mm, #0 coverglass bottoms (MatTek) and overlaid with sterile 50% glycerol. Nodule images were collected on a Leica TCS SPE laser-scanning confocal (model DMI4000B-CS) using a 10X/0.3 NA APO ACS objective and solid-state laser lines for fluorophore excitation at the following settings for each dye: Calcofluor, 405 nm excitation/410-500 nm emission; SYTO9, 488 nm excitation/510-570 nm emission; PI, 532 nm excitation/600-650 nm emission. These images were then processed to enhance brightness and contrast in FIJI(Schindelin et al. 2012; Schneider et al. 2012).

### Nodule diameter and volume measurements

Inoculated *A. afraspera* root nodules were imaging using a high-definition Keyence VHX-600 digital microscope at 20X magnification. For end-point root nodule volume measurements at 40 days post-inoculation, plants were removed from the growth chamber and imaged at RT on paper towels, then discarded. Nodule diameters were measured using the line tool in FIJI and recorded using a custom FIJI macro. For tracking nodule volumes over time, plants were serially removed from their growth chambers and transferred to a plastic dish containing 150 mL of sterile BNM pre-warmed to 28°C. Images of sections of the plant root were collected serially from the hypocotyl to the root tip. Following collection of images, plants were immediately returned to their original growth tubes in the growth chamber. Plastic dishes were sterilized for 10 minutes in 10% bleach, washed three times in sterile-filtered nanopure water, sprayed with 70% ethanol/water, and air-dried before each new plant was imaged. A fresh aliquot of sterile, pre-warmed BNM also was used for each plant. After the time course was completed, images of entire plant root systems were reconstructed by eye for each plant at each time point. For nodules appearing in at least five time points, nodule diameters were measured as described for the end-point measurements and were converted to approximate volumes in R using the equation *V =* 4/3 *πr*^*3*^.

### Nodule growth curve fitting and analysis

All analyses of nodule growth, and corresponding plots, were generated in R. For nodule growth curve fitting, three model equations were used to identify the best fit, as follows:

(1) exponential function:

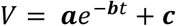

(2) quadratic function:

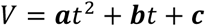

(3) generalized logistic function (expressed as a Richard’s function with a time shift):

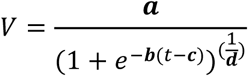

Calculation of the optimal parameter values for each equation (e.g. the values of **a**, **b**, **c**, and **d**) and the standard error for each curve compared to the raw data were performed using the built-in function *nlm*() in R. In some cases, *nlm()* could not produce a best-fit model without specifying initial values for the function parameters. For exponential models, an equation of best fit could be successfully determined without specification of initial values for parameters **a**, **b** and **c**. For quadratic models, initial parameter values were required and were set to **a**=0, **b**=10 and **c**=0 for each nodule plot, after identifying these initial parameter values as broadly optimal based on an initial parameter sweep of −50 to 50 for each plot. For sigmoidal models, no broadly optimal initial values could be identified, so a parameter sweep was performed for each plot with the initial value of **a** set to the maximum observed nodule volume (as **a** describes the upper asymptote of the sigmoidal curve), **b** ranging from 0.1 to 1, **c** ranging from 0 to 10, and **d** ranging from 0.01 to 1.0. In the sigmoidal plots, an initial point of (0,0) was added to the nodule volume time series to improve fitting.

Because the sigmoidal model provided the best fits, extrapolation of nodule growth characteristics was performed on sigmoidal models only. The maximum nodule volume, **V_max_**, is defined as the upper asymptote of the sigmoidal growth curve, *e.g.* **a**. The nodule initiation time, **t_min_**, was defined in three separate ways: the times at which the nodule volume is equal to 0.05, 0.1, or 0.2 mm^3^ (*e.g*. through solving 0.05, 0.1, or 0.2 = **a**/((1+*e*^(-**b**(*t*-**c**))^)^(^1^/**d**)^) for *t*). The maximum nodule growth rate, **dV/d*t***, was defined as the average rate of growth (*e.g.* slope) between the time at which the volume is 10% of **V_max_** and the time at which the volume is 90% of **V_max_**. The time at which each nodule reaches its maximum size, **t_max_**, was approximated as the time at which the volume is 90% of **V_max_**, since the “true” maximum volume is asymptotic to the growth curve and is therefore never fully reached in the model.

### Competition assays

mCherry-tagged Δ*hpnH* and YFP-tagged wild type *B. diazoefficiens* were grown to stationary phase (OD_600_ > 1.4) in 10 mL PSY cultures supplemented with 20 µg/mL (wild type) or 10 µg/mL (Δ*hpnH*) tetracycline; untagged strains were grown in PSY. On the day prior to inoculation, all strains were diluted into 50-150 mL tetracycline-free PSY to reach an OD_600_ of ∼0.8 at the time of inoculation. *A. afraspera* plants were cultivated pre-inoculation in test tubes as described above, with the addition of covering the growth tubes in foil to minimize the production of chlorophyll in the plant roots, which spectrally overlaps with mCherry. At the time of inoculation, all cultures were pelleted at 3250 x g for 30 minutes at RT, washed three times, then resuspended in PSY to a final OD_600_ of 1.0. A 10 mL culture of each strain ratio for inoculation was generated a sterile 15mL Falcon tube; for example, for a 50:50 mixture of mCherry-tagged Δ*hpnH* and YFP-tagged wild type, 5 mL of each strain was combined. These cultures were mixed thoroughly by gentle pipetting, and 1 mL of the mixtures was added to directly to the plant medium for 7-8 plants per strain mixture.

After 45-60 days, plants were harvested. First, plant heights and the number of nodules per plant were recorded. Then, the roots were cut from the stem and images of all nodules for each plant were collected on a high-definition Keyence VHX-600 digital microscope at 20X magnification. These nodules were then cross-sectioned and immediately transferred to Eppendorfs containing 4% paraformaldehyde (Electron Microscopy Sciences) in PBS. Fresh sections were fixed overnight in the dark at 4°C, washed 5X in PBS, and stored in PBS supplemented with 0.1% azide in the dark at 4°C until imaging.

Fixed sections were stained in Calcofluor (all strain combinations), SYTO9 (WT-YFP and WT co-inoculation only) or propidium iodide (mCherry-Δ*hpnH* and Δ*hpnH* co-inoculation only) as described for Live:Dead staining. Imaging was performed as described for Live:Dead staining using a 5X objective. Given the high autofluorescence of these nodules and low mCherry and YFP signal intensities, the following excitation/emission settings were used: Calcofluor, 405 nm excitation/410-460 nm emission; YFP/SYTO9, 488 nm excitation/500-550 nm emisasion; mCherry, 532 nm excitation/600-650 nm emission.

Quantification of nodule statistics (including nodule and infection zone areas, signal intensity of YFP, mCherry, SYTO9 and propidium iodide) was performed on raw images using a custom FIJI macro. Briefly, nodule images were opened at random, infection zones (IZs) and whole nodules were circled by hand and saved as discrete regions of interest (ROIs), and the area and intensity in each channel were measured automatically for all ROIs. These measurements were exported as a text table and various parameters from these measurements were calculated using custom Python scripts, as indicated in the Results. Plots of all parameters and statistical comparisons were generated using custom R scripts.

### Antibiotic treatment of inoculated plants

*afraspera* plants were cultivated as described above and the following antibiotics were added to non-inoculated plants 7 days after rooting in 100 mL BNM growth tubes: kanamycin to 100 µg/mL, streptomycin to 100 µg/mL, tetracycline to 20 µg/mL, kanamycin plus tetracycline, kanamycin plus streptomycin, streptomycin plus tetracycline. Plants were grown in antibiotics under normal plant growth conditions for 14 days, after which plants were visually inspected. Plant heights were also recorded, and the root and shoot systems were separated with a razor blade, transferred into pre-weighed 15 mL Falcon tubes, dried at 50°C for a minimum of 48 hours, then weighed again.

Antibiotic treatments of Δ*hpnH* and wild-type *B. diazoefficiens* were performed by growing antibiotic 5 mL PSY cultures of each strain to stationary phase (OD_600_ >1.4) and diluting strains in fresh PSY to reach an OD_600_ of ∼0.8 at the time of antibiotic treatment – *e.g.* as they would be grown prior to plant inoculation. Cultures were pelleted at 3250 x g for 30 minutes at RT, washed three times, then resuspended in PSY to a final OD_600_ of 1.0. Four 100 µl aliquots of these culture were diluted 1:00 into separate 10 mL BNM cultures in clear glass tubes in plant growth chambers. Kanamycin (at 25, 50, 75, and 100 µg/mL) and streptomycin (at 25, 50, 75, and 100 µg/mL) were added directly to the BNM cultures, and 100 µl samples were taken immediately prior to antibiotic treatment and at 2, 4, 6, 8, and 10 hours post-antibiotic addition. These 100 µl samples were immediately diluted 1:10 in 900 µl and mixed vigorously by repeated pipetting. Vortexing was avoided as we found that this method reduces Δ*hpnH* viability. Ten serial 1:10 dilutions were performed, and three 10 µl samples of each dilution for each strain were spotted and dripped across PSY plates. After 7 days (wild type) or 10 days (Δ*hpnH*), colonies were counted manually and recorded for each dilution exhibiting discrete colonies. Log plots of colony counts over time were generated in R.

Plants were then inoculated with Δ*hpnH* and wild-type *B. diazoefficiens* as described above, and kanamycin and streptomycin were added to Δ*hpnH*-inoculated plants to 50 µg/mL each, and to wild type-inoculated plants to 100 µg/mL at 12 hours and 36 hours and at 2, 2.5, 3, 3.5, 4, 4.5, 5, 6.5, 8.5, 9.5, 10.5, and 12.5 days post-inoculation. Four plants were treated per time point per strain, with an additional four plants each as an untreated control. At 40 dpi, the number of nodules per plant was recorded.

### Bulk motility assays

Swimming motility assays were performed as previously described, with some modifications (Althabegoiti et al. 2008). WT and Δ*hpnH* were grown to turbidity in 5 mL of PSY at 30°C and 250 rpm, diluted to an OD_600_ of 0.02 in 5 mL of fresh PSY, and grown to exponential phase (OD_600_ = 0.3-0.5). Exponential cultures then were diluted to an OD_600_ of 0.06 in fresh PSY and 2 µL of the adjusted cultures into the center of swimming plate containing 0.3% agar/PSY. After inoculation, the plates were wrapped with parafilm to prevent dehydration and incubated in a humidity-controlled environmental chamber (Percival) at 30°C for 10 days total, with daily scans after 5 days. The resulting images were analyzed in FIJI to measure the area of the swimming colony.

### Surface attachment assays

Δ*hpnH* and wild-type *B. diazoefficiens* were grown in 5 mL PSY cultures to stationary phase (OD_600_ >1.4) then diluted in fresh PSY to reach an OD_600_ of ∼0.8 at the time of surface attachment assays. Cultures were pelleted at 3250 x g for 30 minutes at RT, washed twice in the indicated attachment medium, then resuspended in attachment medium to an OD_600_ of 1.0. These cultures were mixed thoroughly by repeated pipetting, and 2 mL samples were added to sterile imaging dishes (30 mm dishes with 20 mm, #1.5 coverglass bottoms; MatTek). Cultures were incubated on imaging dishes *without shaking* at 30°C for two hours. To remove non-adhered cells, imaging dishes were immersed in 50 mL of attachment media in a 100 mL glass beaker on an orbital shaker and shaken gently at RT for 5 minutes; direct application of washing medium to the coverglass surface was avoided, as we found that this creates a shear force sufficient to wash away adhered cells. Imaging dishes were then gently lifted out of the washing medium and imaged with a 100X objective on a Lumascope 720 fluorescence microscope (Etaluma). Forty fields of view were recorded for each strain and media combination. These images were processed in FIJI using the Enhanced Local Contrast (CLAHE) plugin (Heckbert and Karel 1994) and converted into a binary image to determine the area of the imaging window covered with adhered cells. Calculation of the fraction of the surface was performed in Excel and statistical analyses were conducted in R. Areas of the surface containing groups of cells larger than 10 µm^2^ in area were ignored in the calculations, as these likely do not represent true attachment events rather than sedimentation of larger cell clumps. BNM used for attachment assays was prepared as described above, with the addition of 1.0 g/mL arabinose. Because BNM contains salt crystals that can sediment onto coverglass and occlude or obscure adhered cells, this medium was passed through a 2 µm filter (Millipore) prior to the attachment experiments.

### Single-cell motility assays and analysis

B. diazoefficiens wild-type and ΔhpnH were grown in 12.5 ml PSY medium at 30°C and 200 rpm to an OD_600_ = 0.6-0.8 from an AG medium plate culture. Then, a 1:10 dilution of cell culture was subcultured in PSY medium to a final volume of 12.5 ml and regrown to an OD_600_ of ∼0.6. Two aliquots of 750 µL were sampled from the regrowth culture and pelleted at 3500 x g for 20 min (wild-type) or for 30 min (ΔhpnH) at RT. The supernatant was removed, and one pellet was resuspended in 500 µL PSY and the other in 500 µL BNM medium. Because BNM contains salt crystals that can sediment onto coverglass and occlude or obscure adhered cells, this medium was passed through a 2 µm filter (Millipore) prior to usage for these experiments. The two medium conditions were then incubated for 2.5 hrs (wild-type) or for 3.5 hrs (ΔhpnH) at 30°C; given the difference in growth time ΔhpnH incubated for longer. Right before imaging, each culture was diluted at a 1:10 ratio with its respective medium. The bacteria were then injected into a sterile flow cell (ibidi sticky-Slide VI0.4 with a glass coverslip). The flow cell was attached to a heating stage set to 30°C.

The imaging protocol involved high-speed bright-field imaging for 5 min at a single XYZ location per experimental repeat. High speed bright-field recordings used a Phantom V12.1 high speed camera (Vision Research); images were taken with a 5 ms exposure at 200 fps and a resolution of 512×512 pixels (0.1 µm/pixel). This protocol was performed on an Olympus IX83 microscope equipped with a 100× oil objective, a 2× multiplier lens, and a Zero Drift Correction autofocus system. The recorded movies were extracted into single frames from the.cine files using PCC 2.8 (Phantom Software). Image processing and cell tracking algorithms are adapted from previous work (Lee et al. 2018) and written in MATLAB R2015a (Mathworks).

We identified cells swimming near the surface as cells with a trajectory radius of gyration greater than 2.5 µm and a mean-squared displacement (MSD) slope greater than 1.5. Setting a minimum radius of gyration selects for cells with a minimum net translation on the across the surface, while a minimum MSD slope threshold ensured the cells are moving super-diffusively (MSD slope ≅ 1, diffusive motion; MSD slope ≅ 2, super-diffusive motion). For each tracked cell, the mean-speed, **v**, was calculated by averaging a moving window, **w**, of the displacement over the cell’s full trajectory, using the following equation:

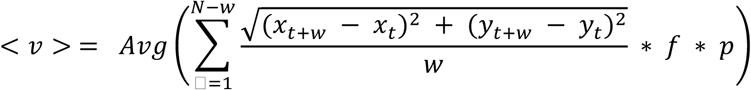

where **N** is the total number of points in the trajectory, **f** is the acquisition frame rate, and **p** is the pixel resolution. Here we set a window size, w= 40 frames. All analysis and visualizations from these experiments where done using MATLAB R2015a (Mathworks).

## Supporting information

## Acknowledgements

This work was supported by grants from the HHMI (D.K.N.), NASA (NNX12AD93G, D.K.N.), Jane Coffin Childs Memorial Fund (B.J.B.), NIH (K99GM126141, B.J.B.), and Army Research Office (W911NF-18-1-0254, GW) and predoctoral fellowships from NSF (E.T.) and the Ford Foundation (J.d.A.). We thank Dr. Eric Giraud for his generous gift of *A. afraspera* seeds and training on *Aeschynomene* symbioses and Drs. Hans-Martin Fischer and Raphael Ledermann for plasmids and technical advice for the genetic transformation of *B. diazoefficiens*. Dr. Nathan Dalleska of the Environmental Analysis Center at Caltech was instrumental in providing training and support for GC-MS analysis of acetylene reduction. We are grateful to Dr. Gargi Kulkarni and other members of the Newman lab, as well as Drs. Elliot Meyerowitz and Rob Phillips, for their collegiality and thoughtful discussions about this work. We are indebted to Ms. Shannon Park and Ms. Kristy Nguyen for providing the administrative assistance that allows us to focus on our research.

## Supplemental Data

**Table S1**. Motile cell counts and mean swimming speeds for wild-type and Δ*hpnH B. diazoefficiens*.

**Figure S1.**
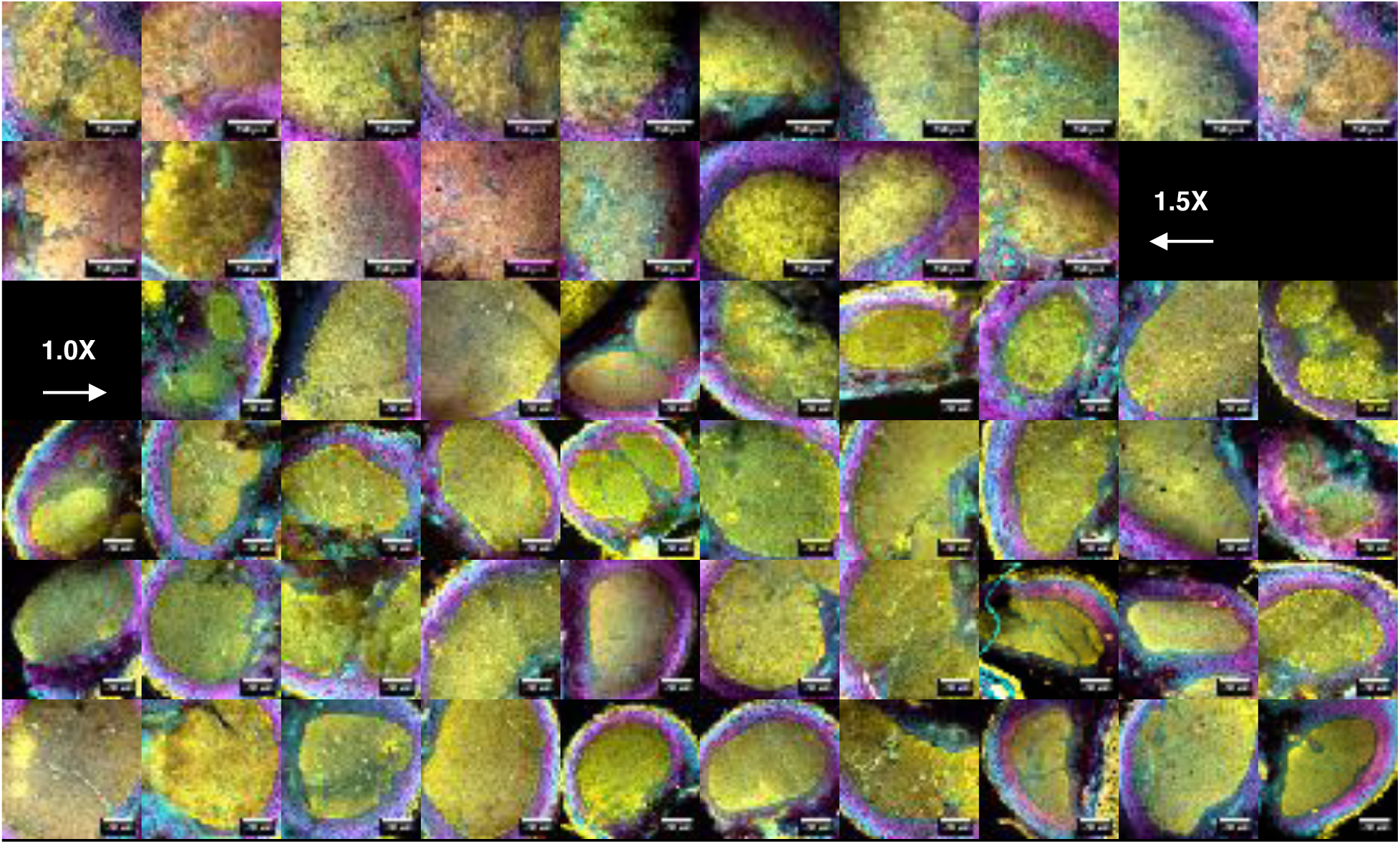
Confocal images of cross-sections of wild type-infected *A. afraspera* nodules at 24 dpi illustrating plant cell walls (Calcofluor, cyan), live bacteria (SYTO9, yellow) and membrane-compromised bacteria and plant nuclei (propidium iodide, magenta). Nodules were collected from 3 plants.

**Figure S2.**
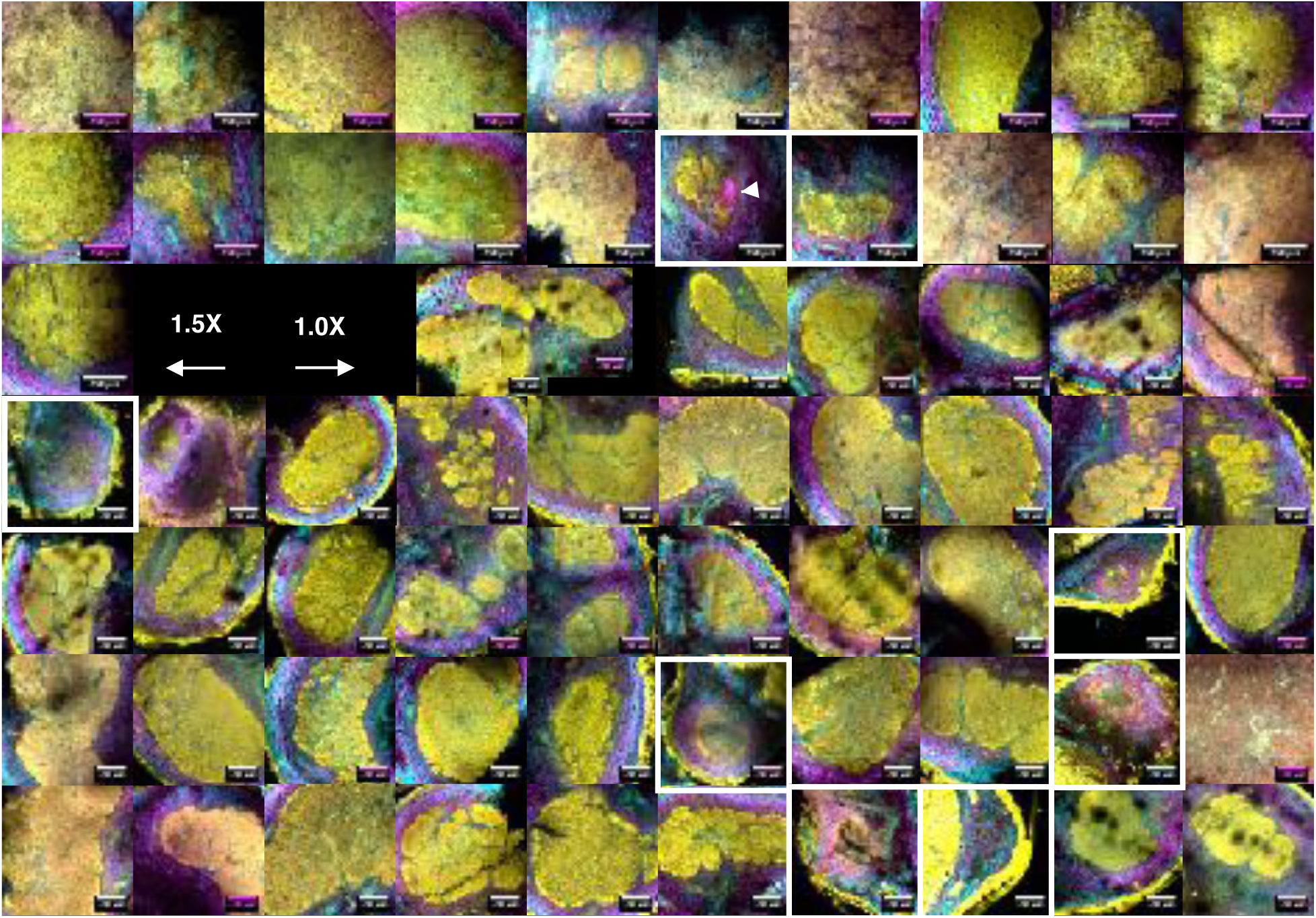
Confocal images of cross-sections of Δ*hpnH*-infected *A. afraspera* nodules at 24 dpi illustrating plant cell walls (Calcofluor, cyan), live bacteria (SYTO9, yellow) and dead bacteria and plant nuclei (propidium iodide, magenta). Nodules were collected from 3 plants. White boxes highlight small nodules. White arrow indicates a likely plant defense reaction.

**Figure S3.**
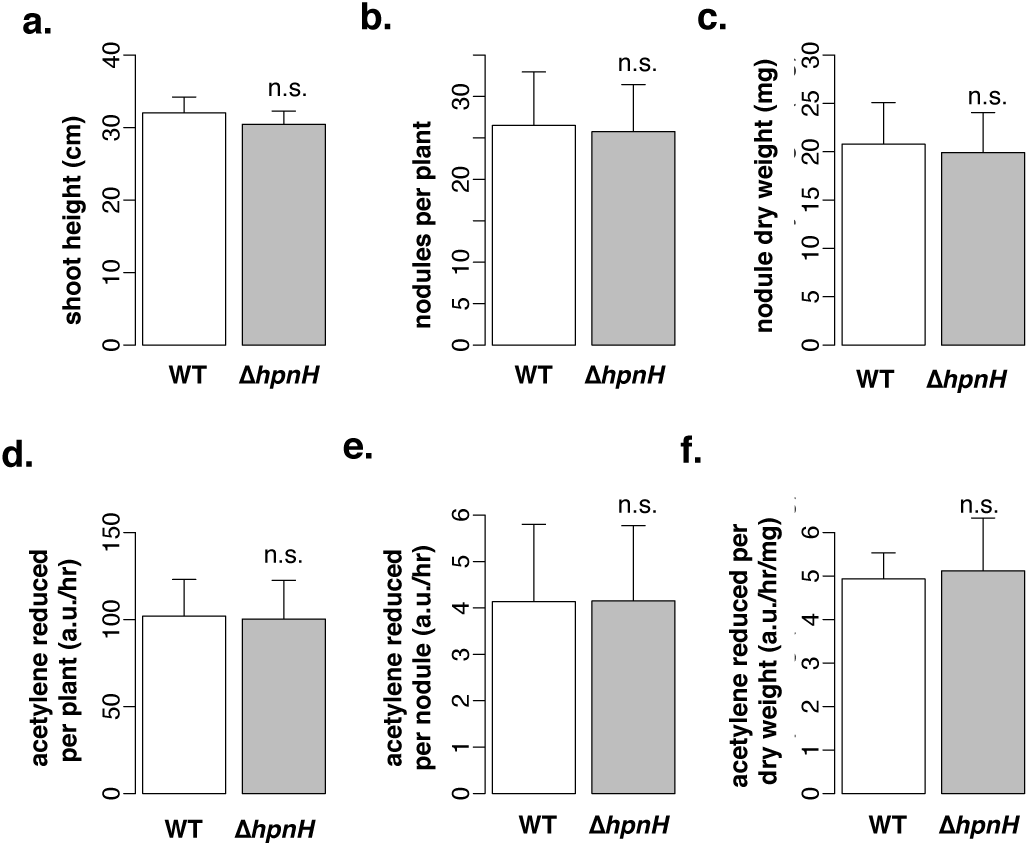
Average (**a**) shoot height, (**b**) nodules per plant, (**c**) nodule dry weight per plant, (**d**) acetylene reduction per plant, (**e**) acetylene reduction per nodule, and (**f**) acetylene reduction per nodule dry weight for *A. asfrapera* inoculated with wild-type or Δ*hpnH* at 40 dpi. N=4 plants per bar; error bars represent one standard deviation. Results of two-tailed t-tests between wild type and Δ*hpnH* are denoted as follows: n.s., p>0.05.

**Figure S4.**
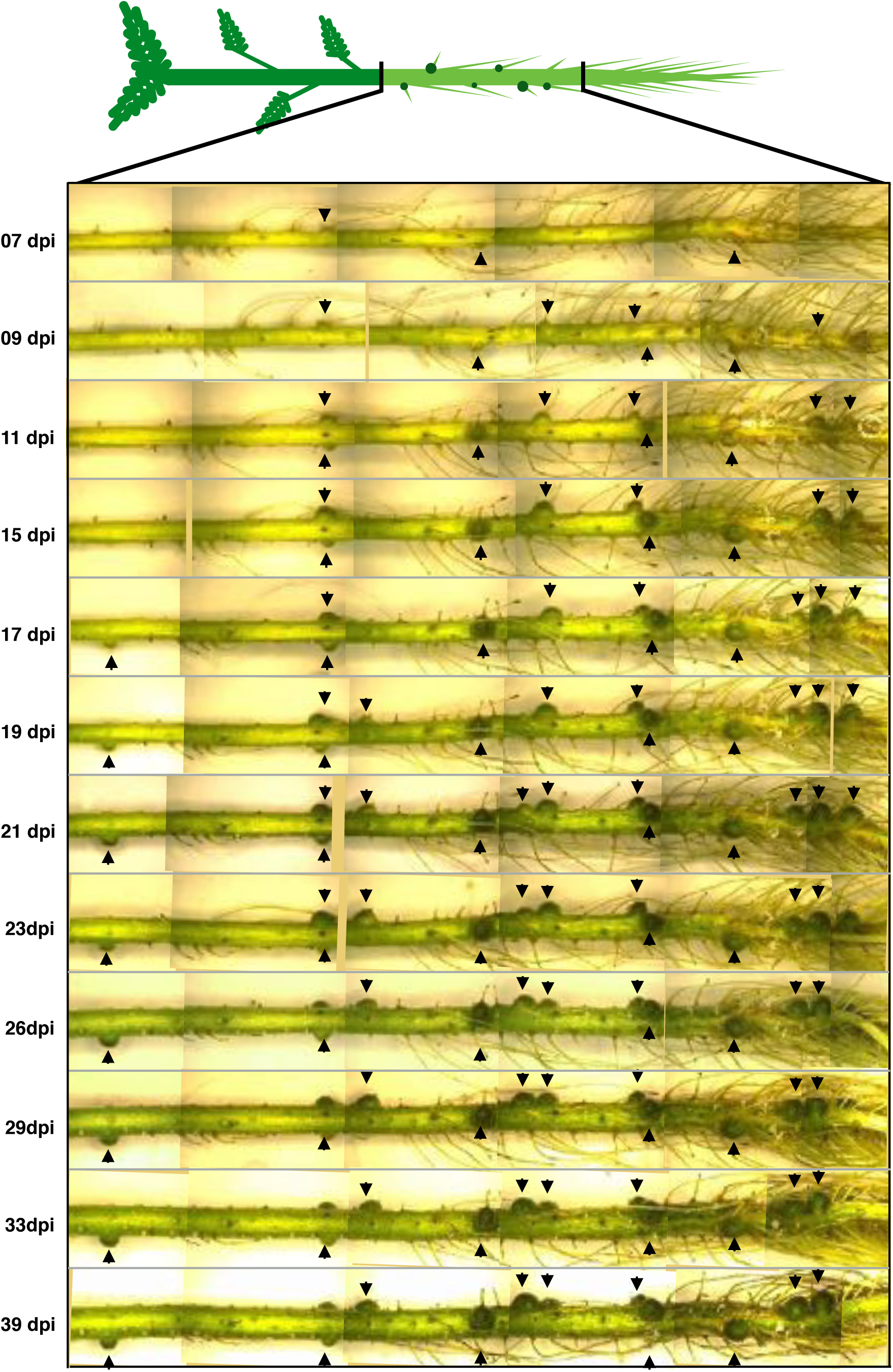
Reconstructed images of the root system of a wild type-infected *A. afraspera* plant. Nodules fully visible in at least five time points are indicated with black arrowheads.

**Figure S5.**
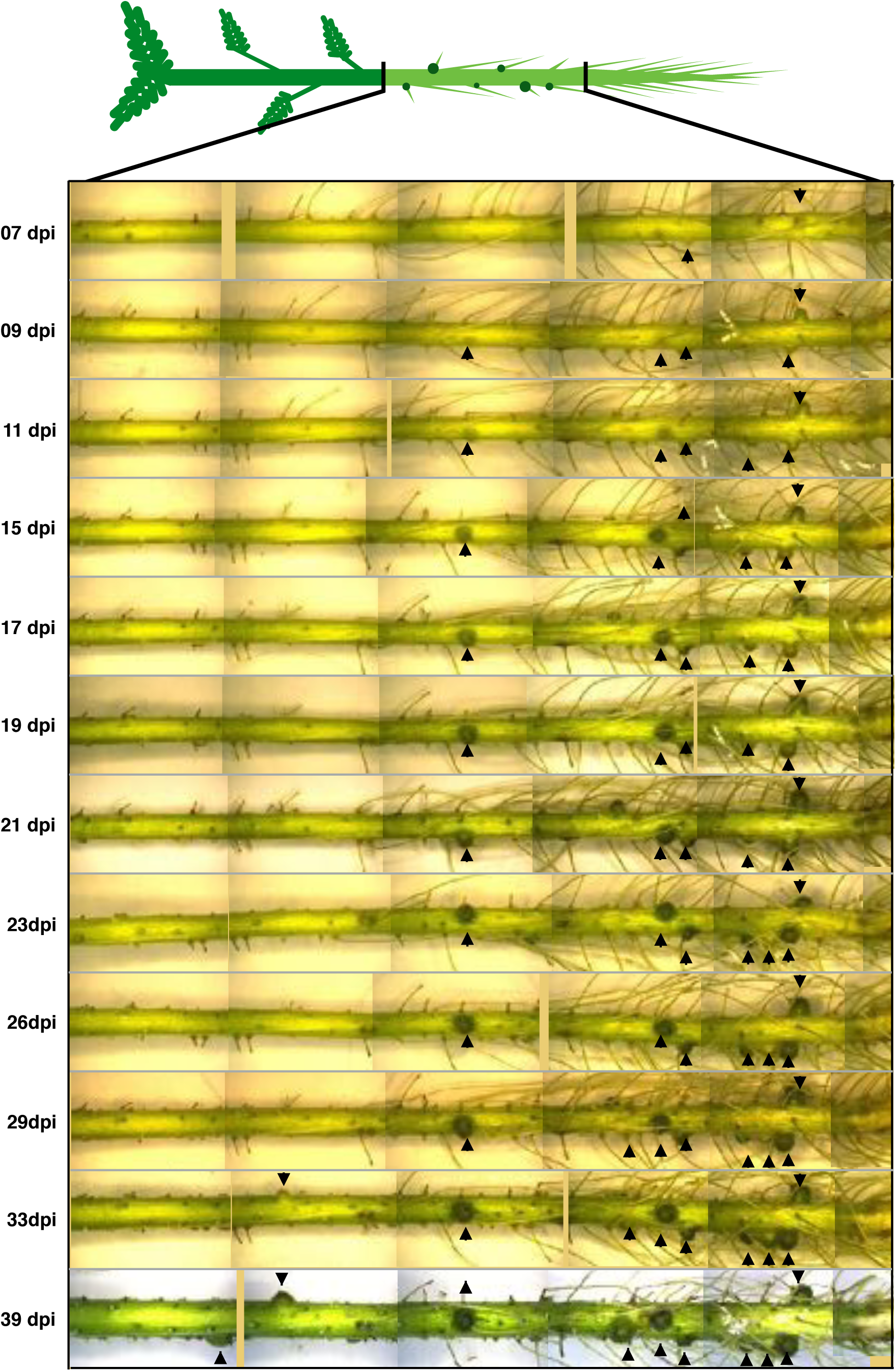
Reconstructed images of the root system of a Δ*hpnH*-infected *A. afraspera* plant. Nodules fully visible in at least five time points are indicated with black arrowheads.

**Figure S6.**
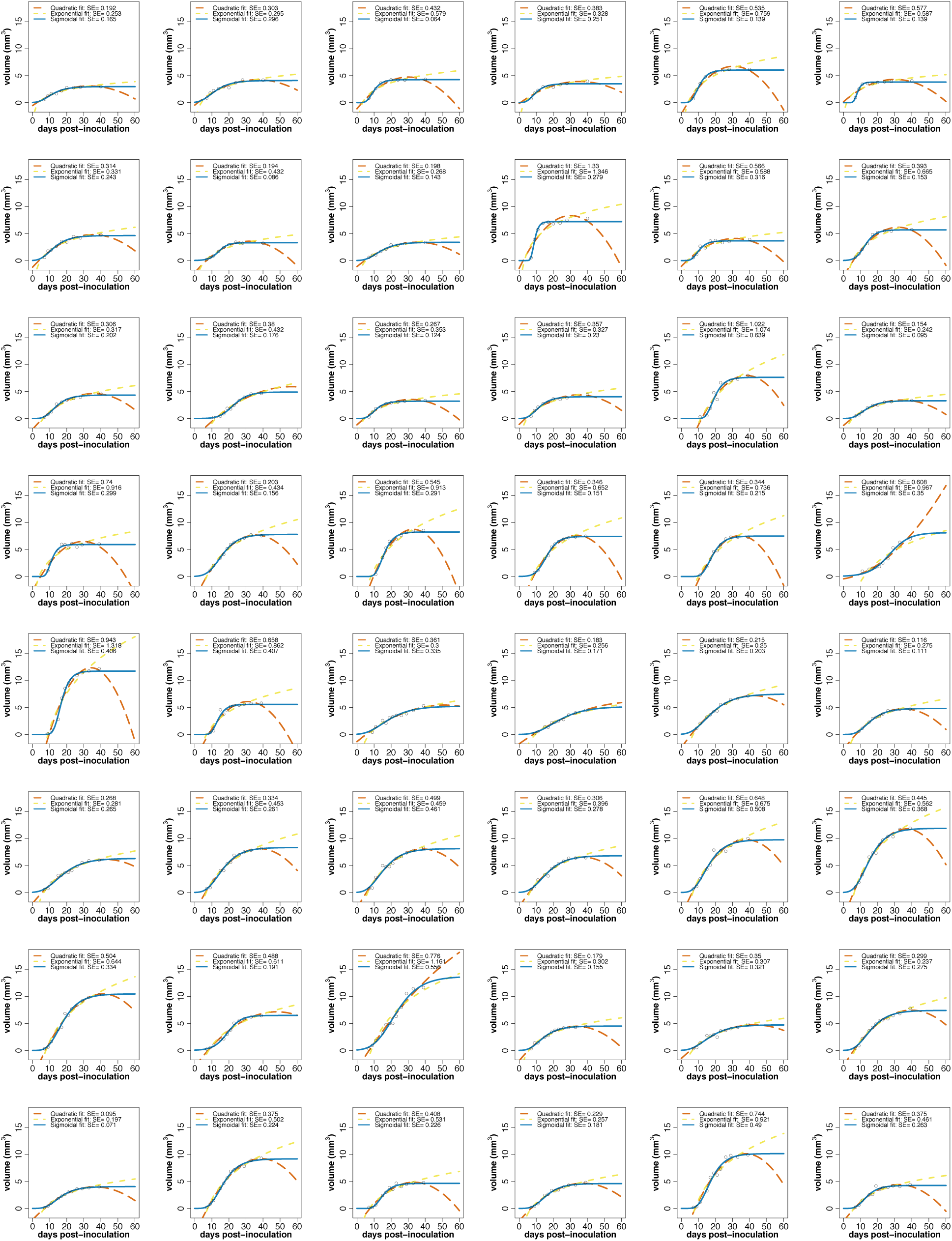
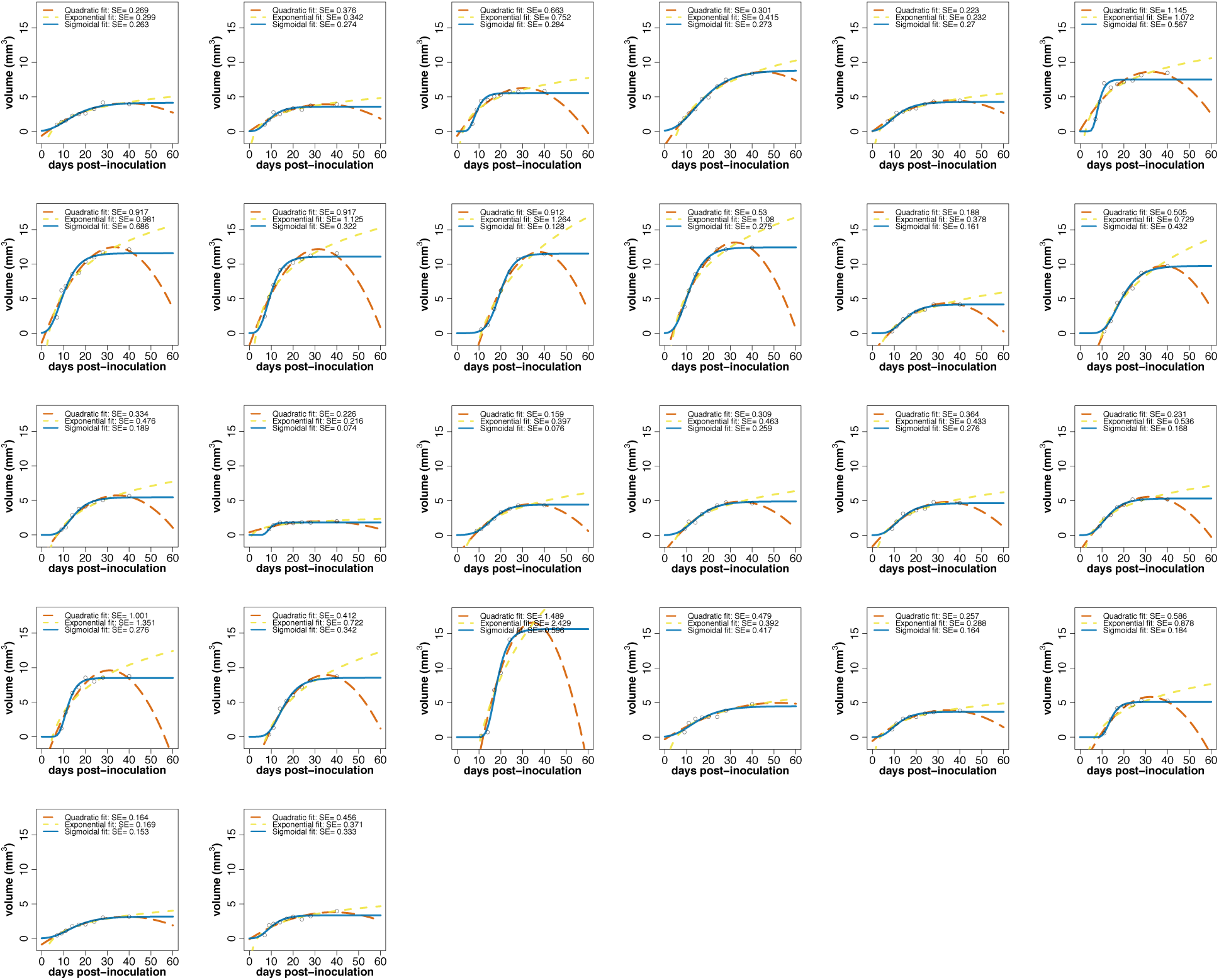
Nodule growth plots for all 74 wild type-infected nodules fit with quadratic (orange; long dashed lines), exponential (yellow; short dashed lines), or sigmoidal (blue; solid lines) models. Standard errors (SE) for each model are shown.

**Figure S7.**
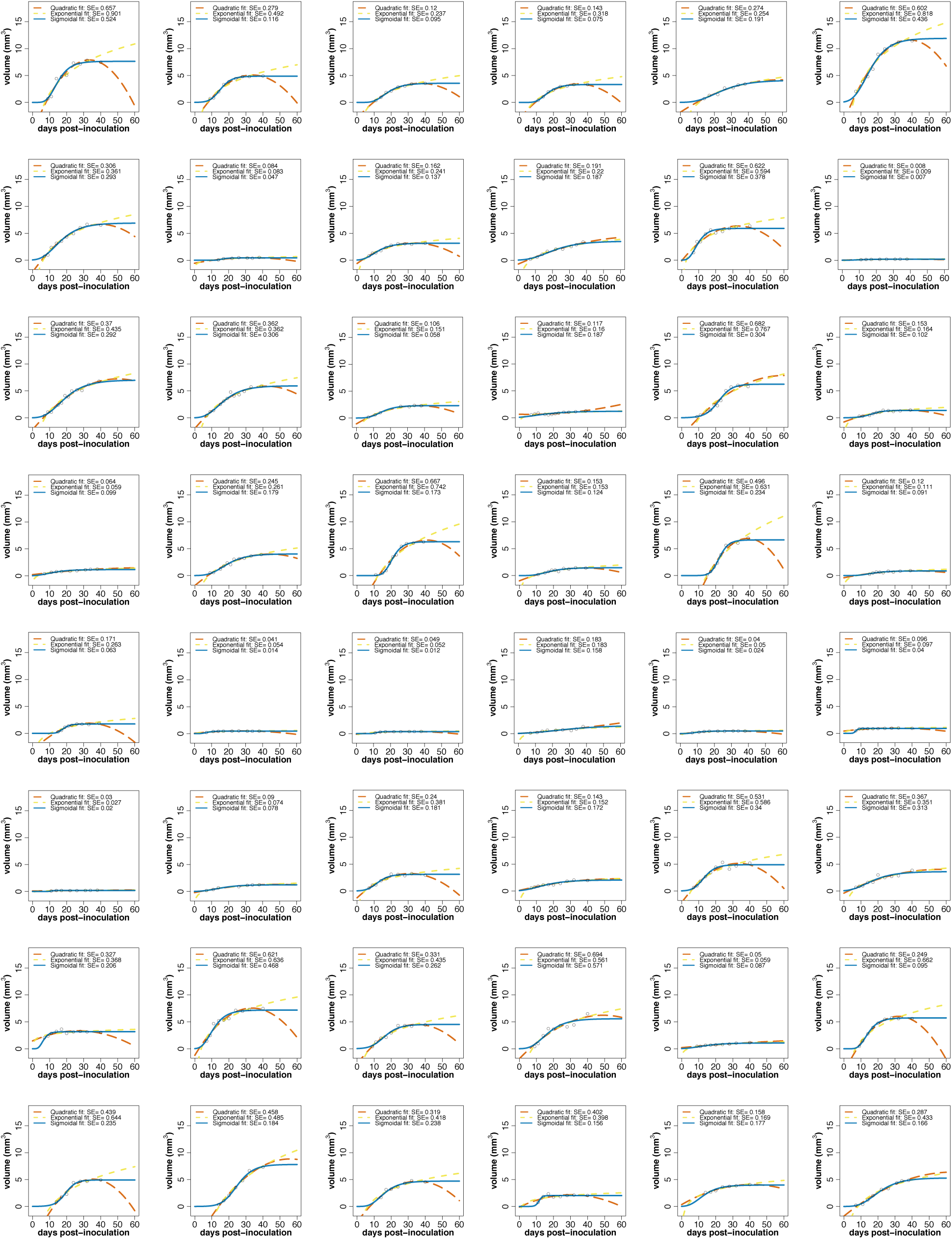
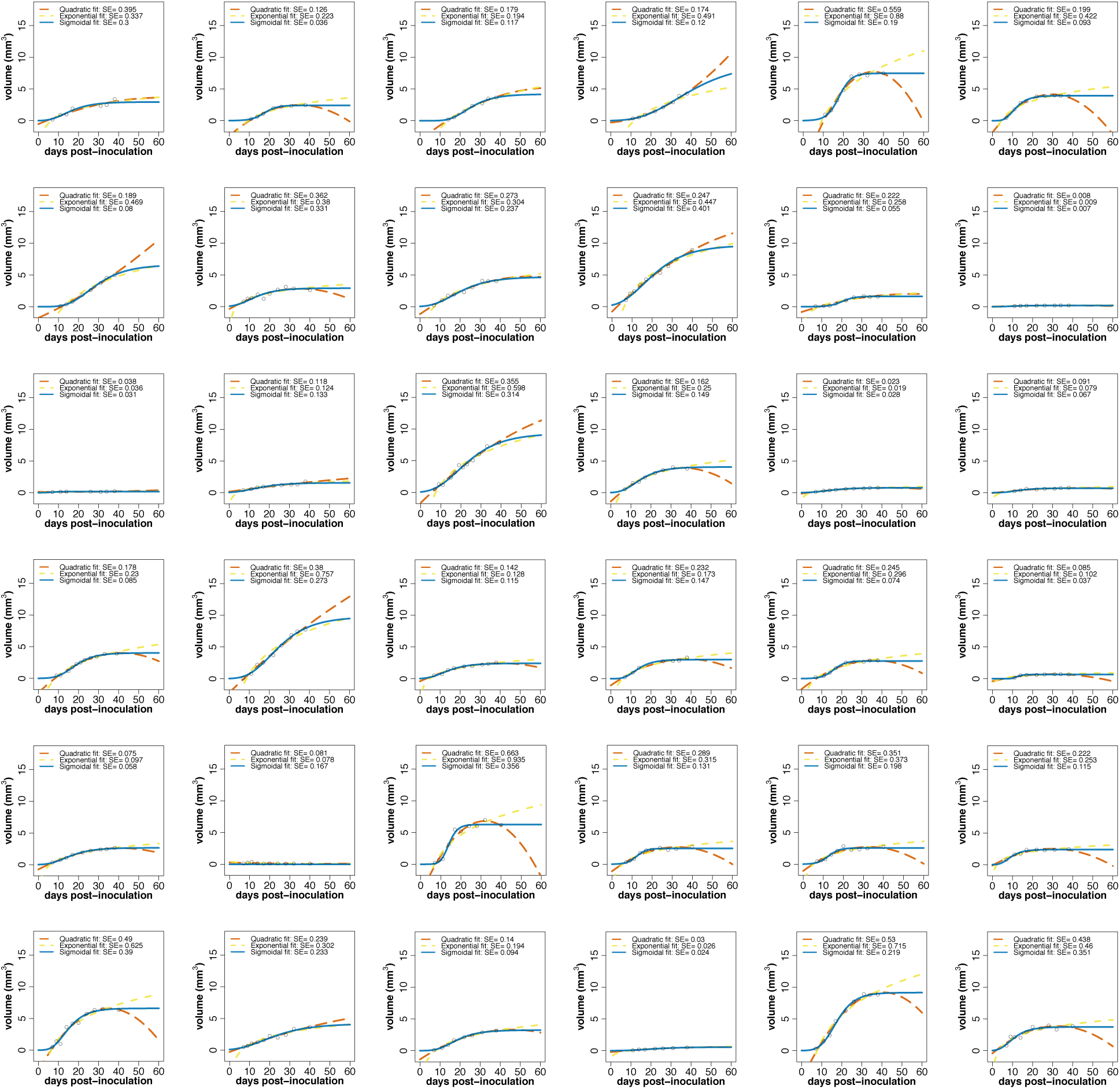
Nodule growth plots for all 84 Δ*hpnH*-infected nodules fit with quadratic (orange; long dashed lines), exponential (yellow; short dashed lines), or sigmoidal (blue; solid lines) models. Standard errors (SE) for each model are shown.

**Figure S8.**
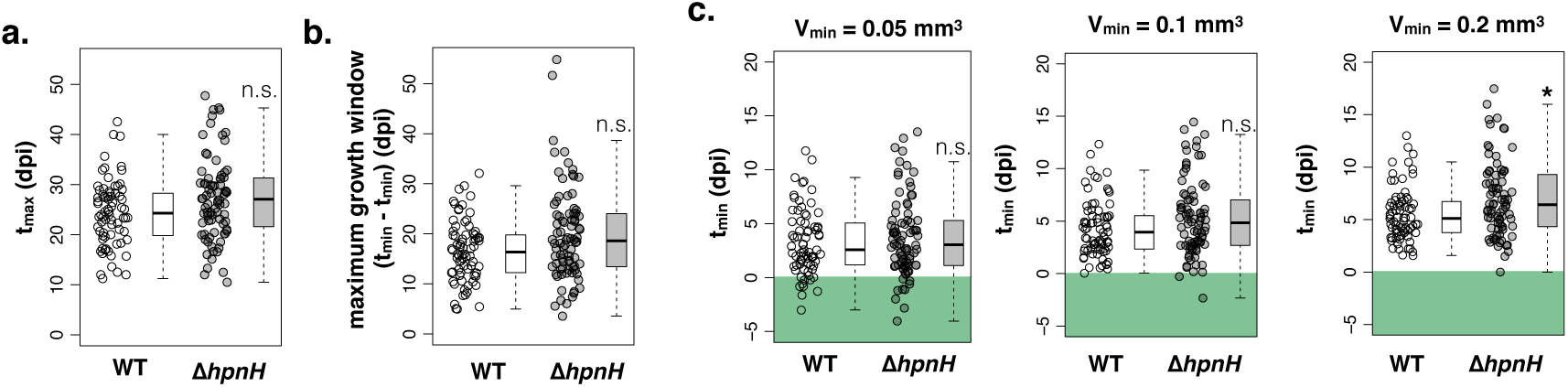
(**a**) Jitter and box plots of **t_max_** values for all wild type- and Δ*hpnH*-infected nodules. (**b**) Jitter and box plots of maximum growth windows for all wild type- and Δ*hpnH*-infected nodules. (**c**) Jitter and box plots of **t_min_** values (as determined by extrapolation using sigmoidal fits of nodule growth curves) for all wild type- and Δ*hpnH*-infected nodules, in which **V_min_** is defined as 0.05 mm^3^, 0.1 mm^3^, 0.2 mm^3^. Green shading highlights negative **t_min_** values. Results of KS-tests between wild-type and Δ*hpnH* nodules are denoted as follows: *, p<0.05; n.s., p>0.05.

**Figure S9.**
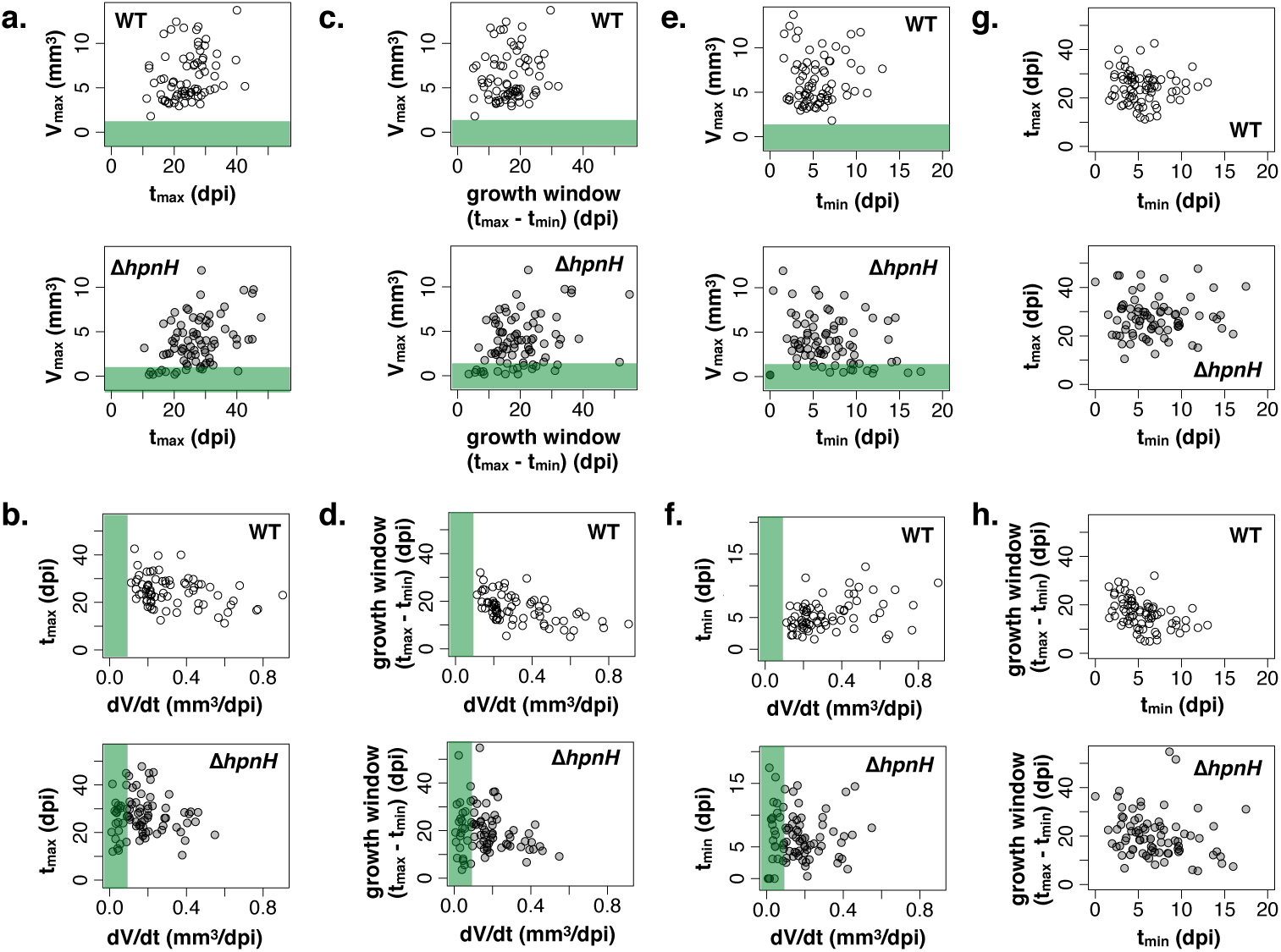
(**a-b**) Scatter plots of **t_max_** vs. (**a**) **dV/dt** and (**b**) **V_max_** for all wild type- (open circles) and Δ*hpnH*- (grey circles) infected nodules. Green regions highlight values below what is observed for wild type. (**c-d**) Scatter plots of maximum growth windows vs. (**c**) **dV/dt** and (**d**) **V_max_**. (**e-f**) Scatter plots of **t_min_** vs. (**c**) **dV/dt** and (**d**) **V_max_**. (**g-h**) Scatter plots of **t_min_** vs. (**a**) **t_max_** and (**b**) maximum growth windows.

**Figure S10.**
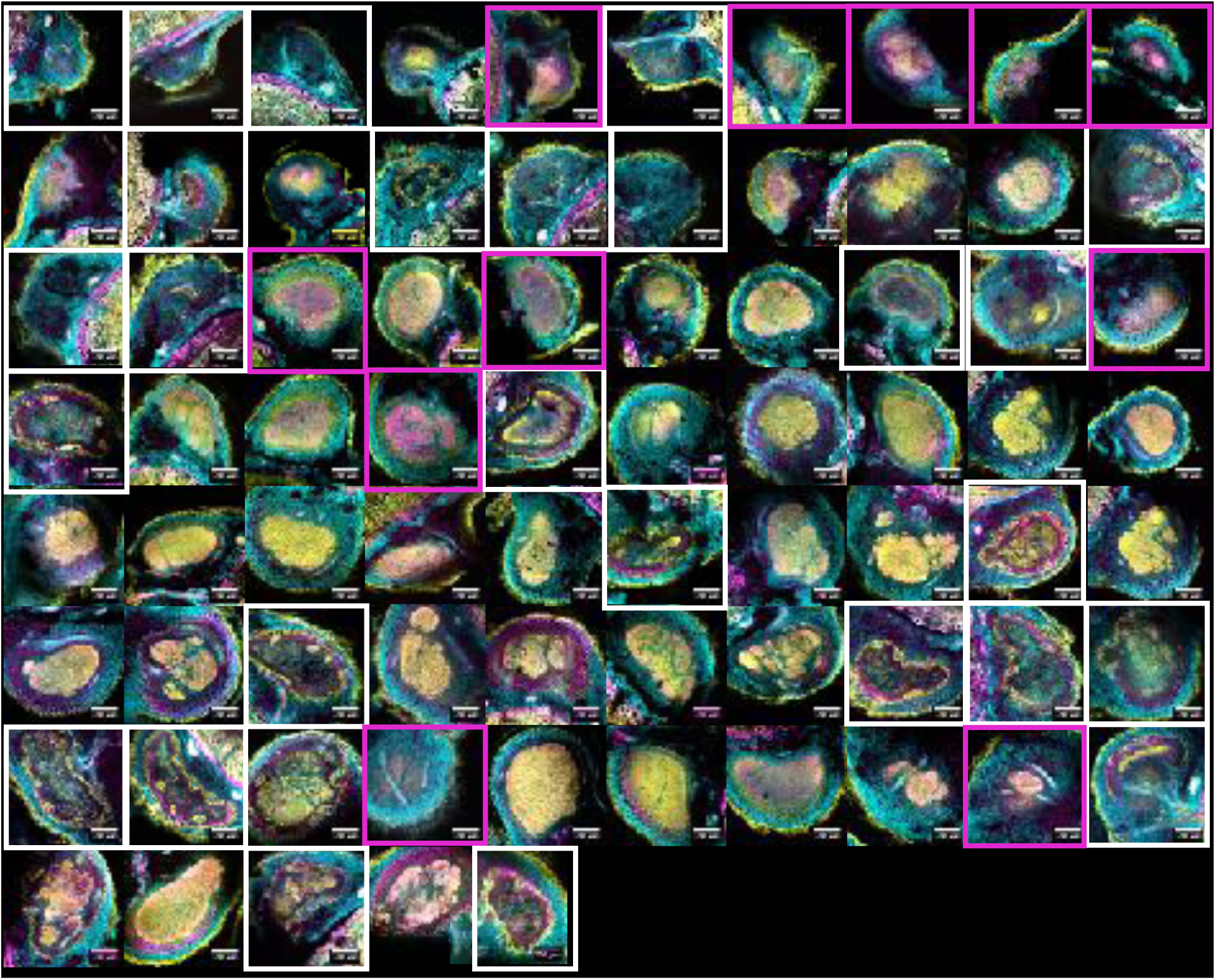
Confocal sections of small (<0.5 mm radius) Δ*hpnH*-infected nodules harvested at 40 dpi. Sections were stained with Calcofluor (cyan), SYTO9 (yellow), and propidium iodide (magenta). N=74 nodules harvested from 5 plants. White boxes highlight under-infected nodules. Magenta boxes indicate nodules primarily containing membrane-compromised cells.

**Figure S11.**
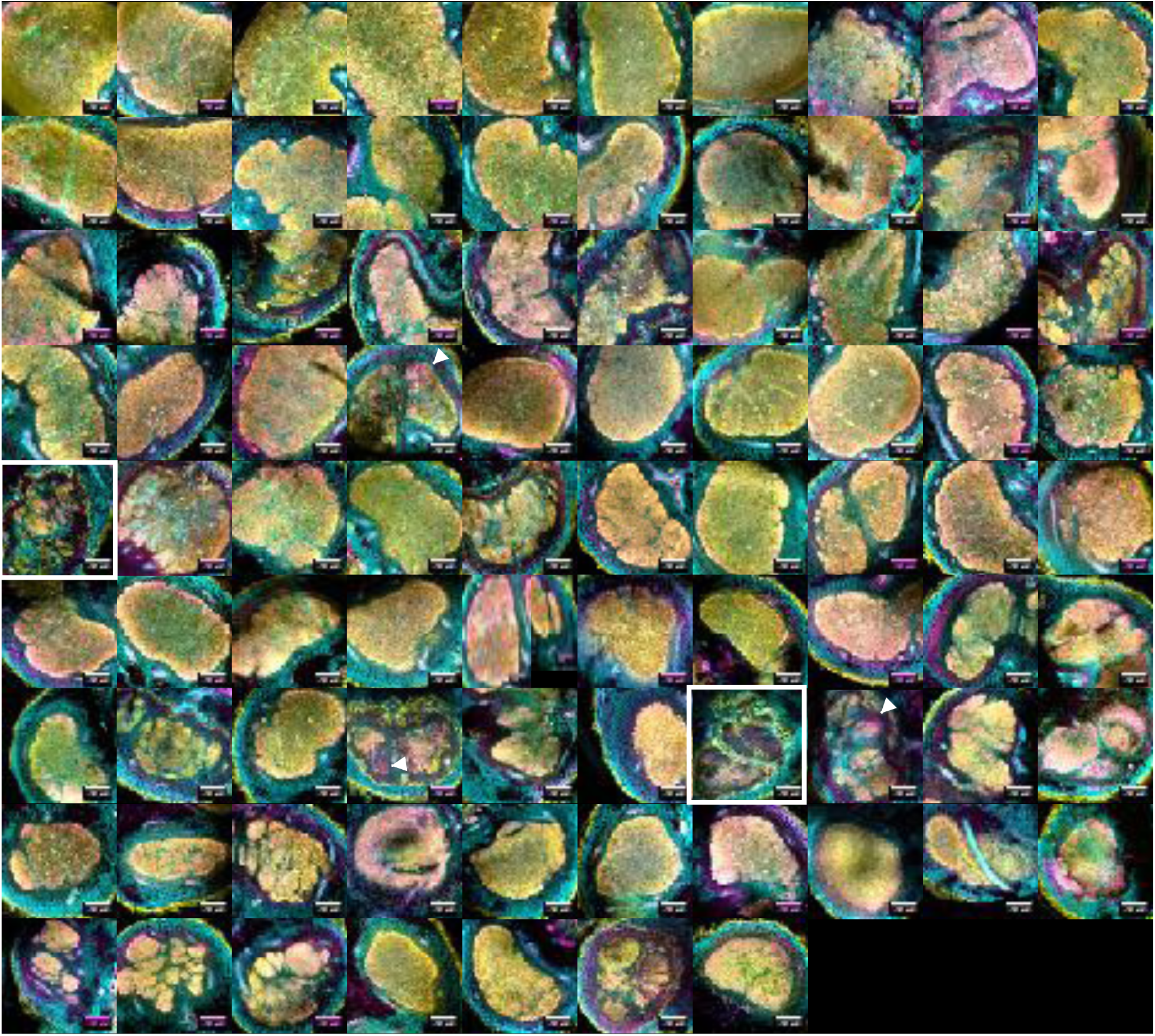
Confocal sections of large (>0.5 mm radius) Δ*hpnH*-infected nodules harvested at 40 dpi. Sections were stained with Calcofluor (cyan), SYTO9 (yellow), and propidium iodide (magenta). N=87 nodules harvested from 5 plants. White boxes highlight under-infected nodules.

**Figure S12.**
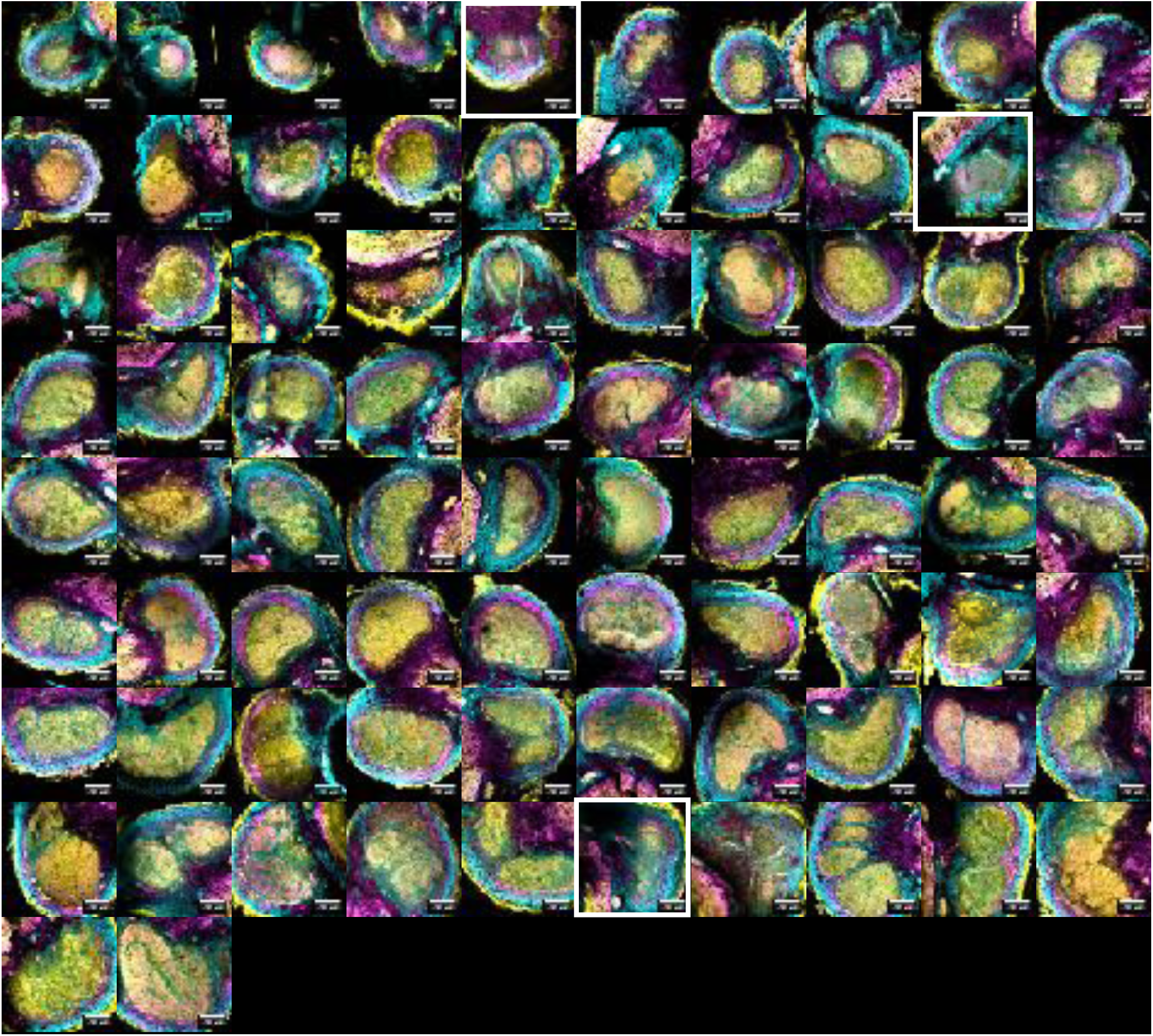
Confocal sections of small (<0.5 mm radius) wild type-infected nodules harvested at 10 dpi. Sections were stained with Calcofluor (cyan), SYTO9 (yellow), and propidium iodide (magenta). N=80 nodules harvested from 5 plants. White boxes highlight under-infected nodules.

**Figure S13.**
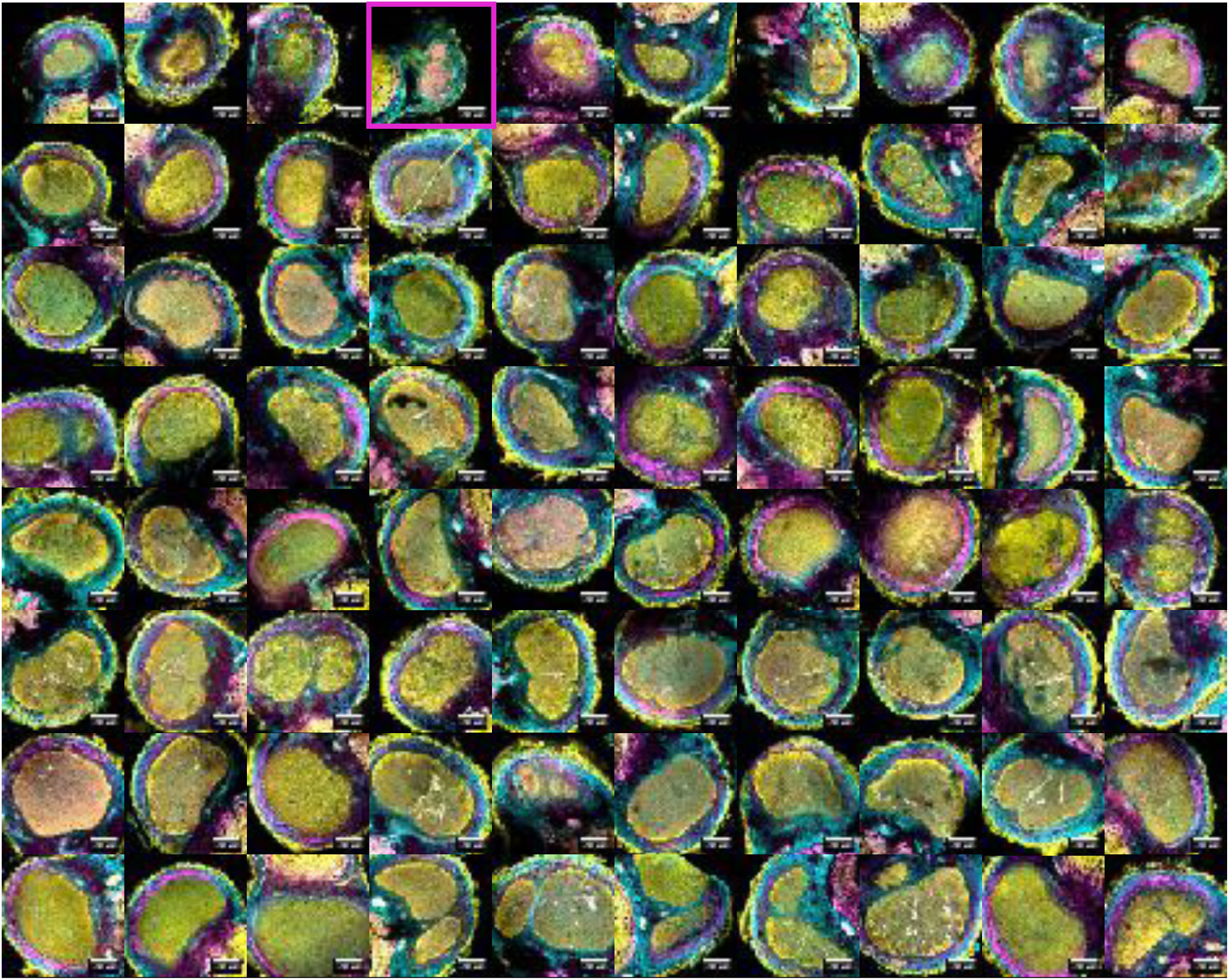
Confocal sections of small (<0.5 mm radius) wild type-infected nodules harvested at 25 dpi. Sections were stained with Calcofluor (cyan), SYTO9 (yellow), and propidium iodide (magenta). N=82 nodules harvested from 5 plants. Magenta boxes indicate nodules primarily containing membrane-compromised cells.

**Figure S14.**
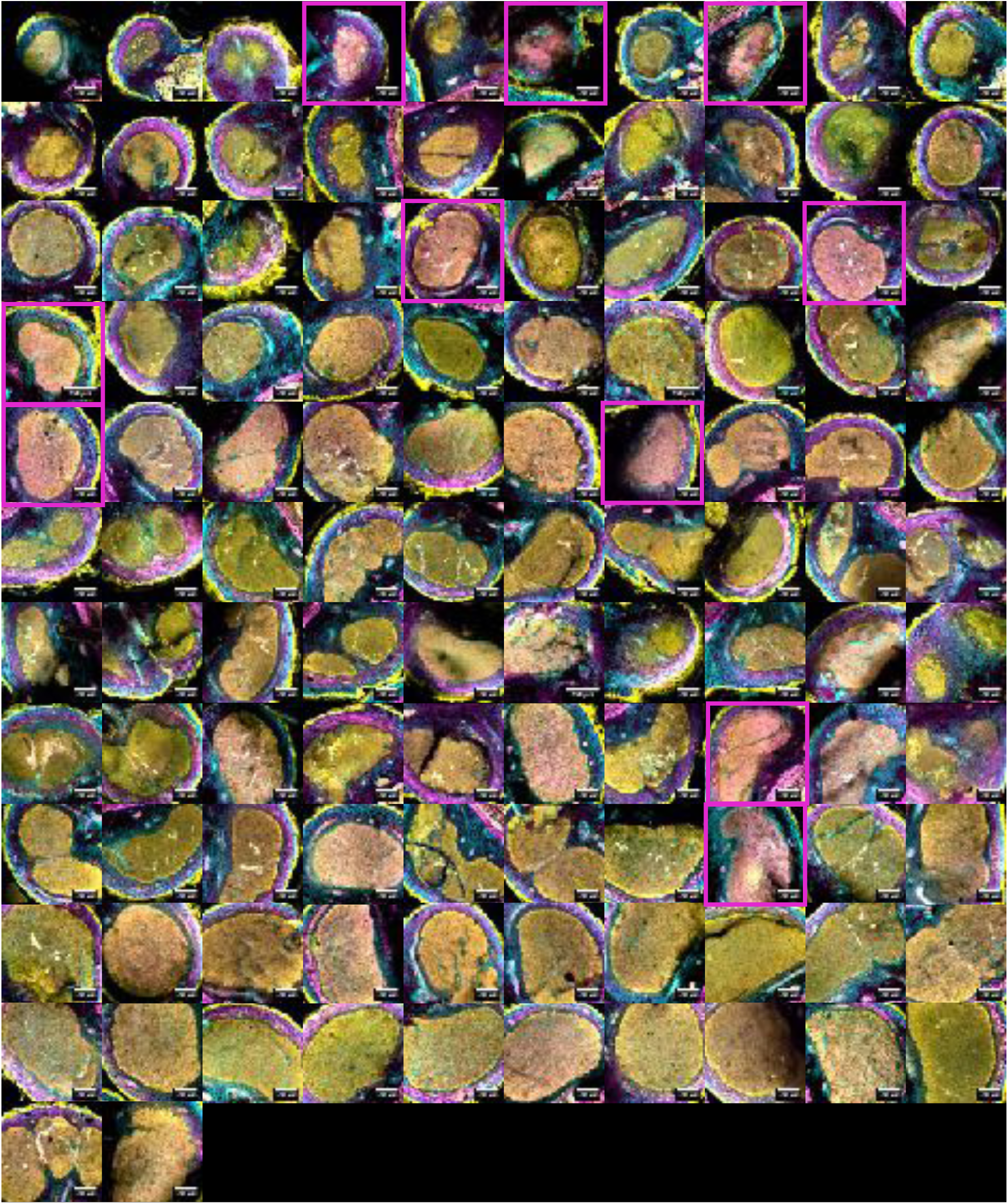
Confocal sections of wild type-infected nodules harvested at 40 dpi. Sections were stained with Calcofluor (cyan), SYTO9 (yellow), and propidium iodide (magenta). N=117 nodules harvested from 5 plants. Magenta boxes indicate nodules primarily containing membrane-compromised cells.

**Figure S15.**
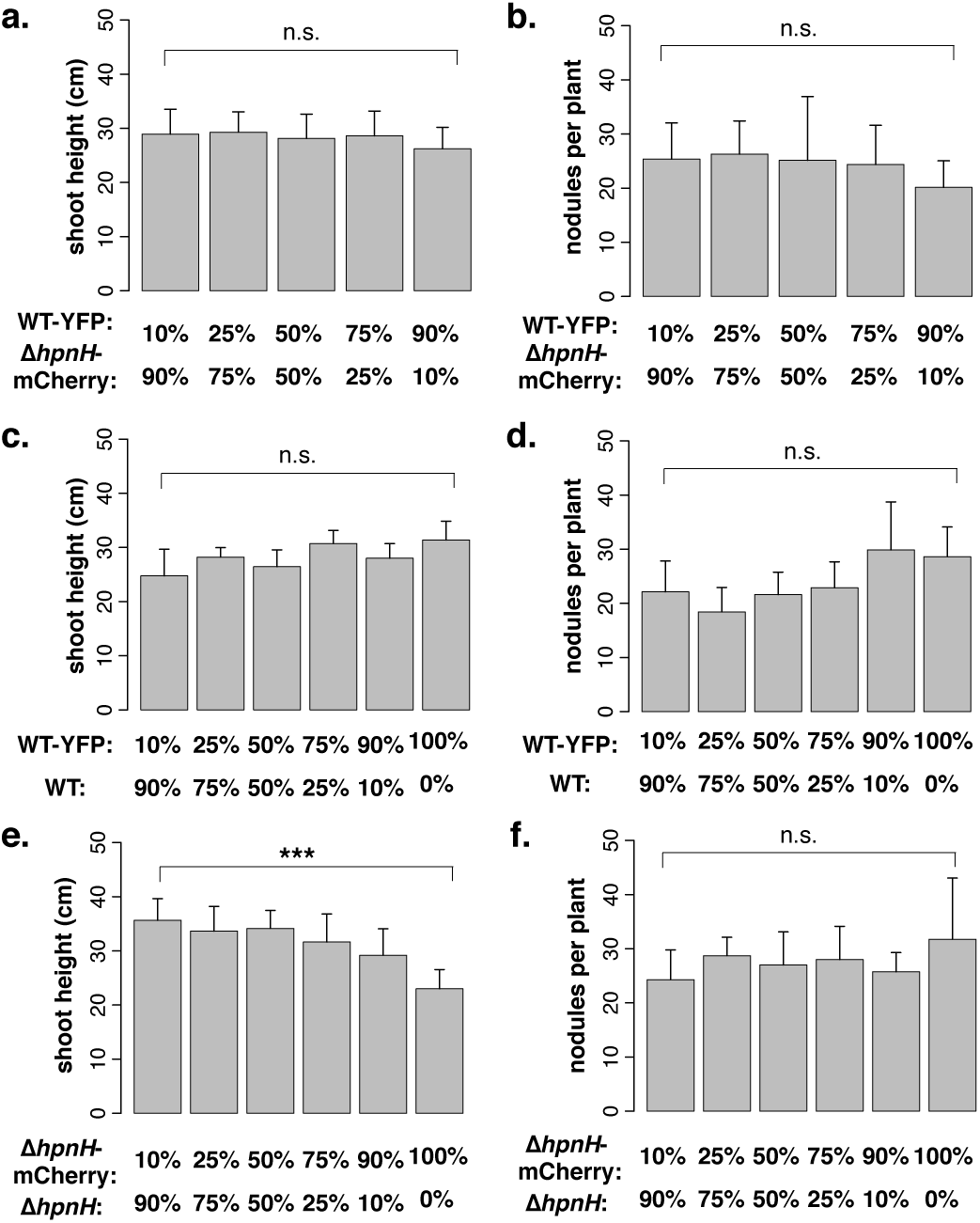
Average shoot height (**a**) and number of nodules (**b**) for plants co-inoculated with Δ*hpnH*- mCherry and WT-YFP strains, recorded at 45 dpi. Average shoot height (**c**) and number of nodules (**d**) for plants co-inoculated with WT and WT-YFP strains, recorded at 40 dpi. Average shoot height (**e**) and number of nodules (**f**) for plants co-inoculated with Δ*hpnH* and Δ*hpnH*-mCherry strains, recorded at 50 dpi. N=7-8 plants per bar for all panels. Error bars represent one standard deviation. Results of two-tailed t-tests are denoted as follows: n.s., p>0.05; ***, p<0.0001.

**Figure S16.**
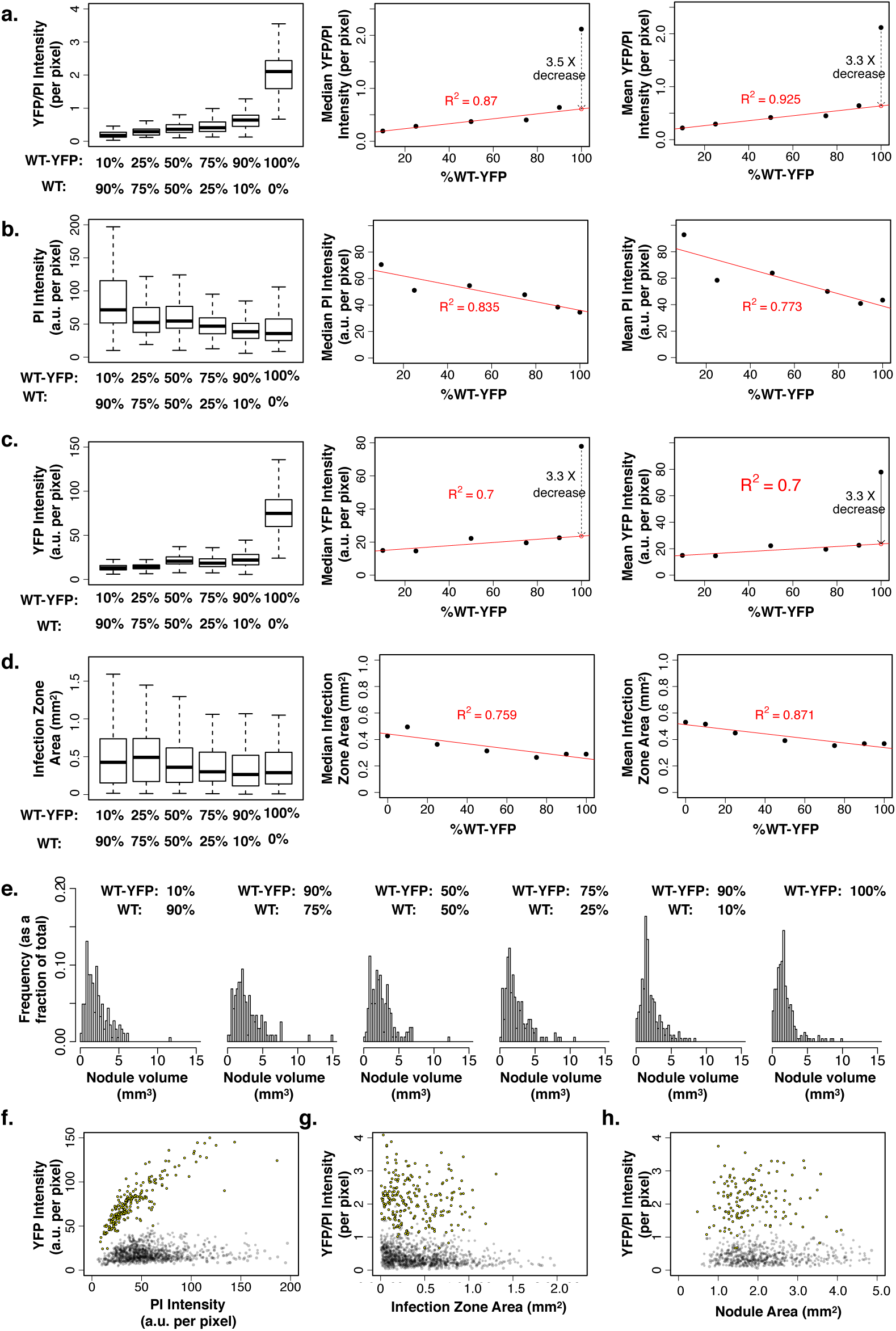
(**a**-**d**) Intensity ratio of YFP to mCherry (**a**), mCherry intensity (**b**), and YFP intensity (**c**) per pixel within infection zones of nodules co-inoculated with Δ*hpnH*-mCherry and WT-YFP strains. (**d**) Cross-sectional area of infection zones of nodules co-inoculated with Δ*hpnH*-mCherry and WT-YFP strains. For (**a**-**d**), N=132, 125, 143, 143 and 110 nodules for 10%, 25%, 50%, 75% and 90% WT-YFP strain mixtures, respectively, which were sectioned and fixed between 45-50 dpi. (**e**) Nodule volume distributions from plants co-inoculated with Δ*hpnH*-mCherry and WT-YFP strains at 45 dpi. Sample sizes are N = 251, 200, 227, 204, and 149 nodules pooled from N = 8, 7, 7, 8, and 7 plants for the 10%, 25%, 50%, 75% and 90% WT-YFP strain mixtures, respectively. (**f**) Scatter plots of mCherry vs. YFP intensities per pixel within infection zones of nodules co-inoculated with Δ*hpnH*-mCherry and WT-YFP strains. (**g**-**h**) Scatter plots of YFP/mCherry intensity ratios per pixel in infection zones vs. infection zone (**g**) and nodule (**h**) cross-section areas for nodules co-inoculated with Δ*hpnH*-mCherry and WT-YFP. Scatter plots contain data pooled from all ratios.

**Figure S17.**
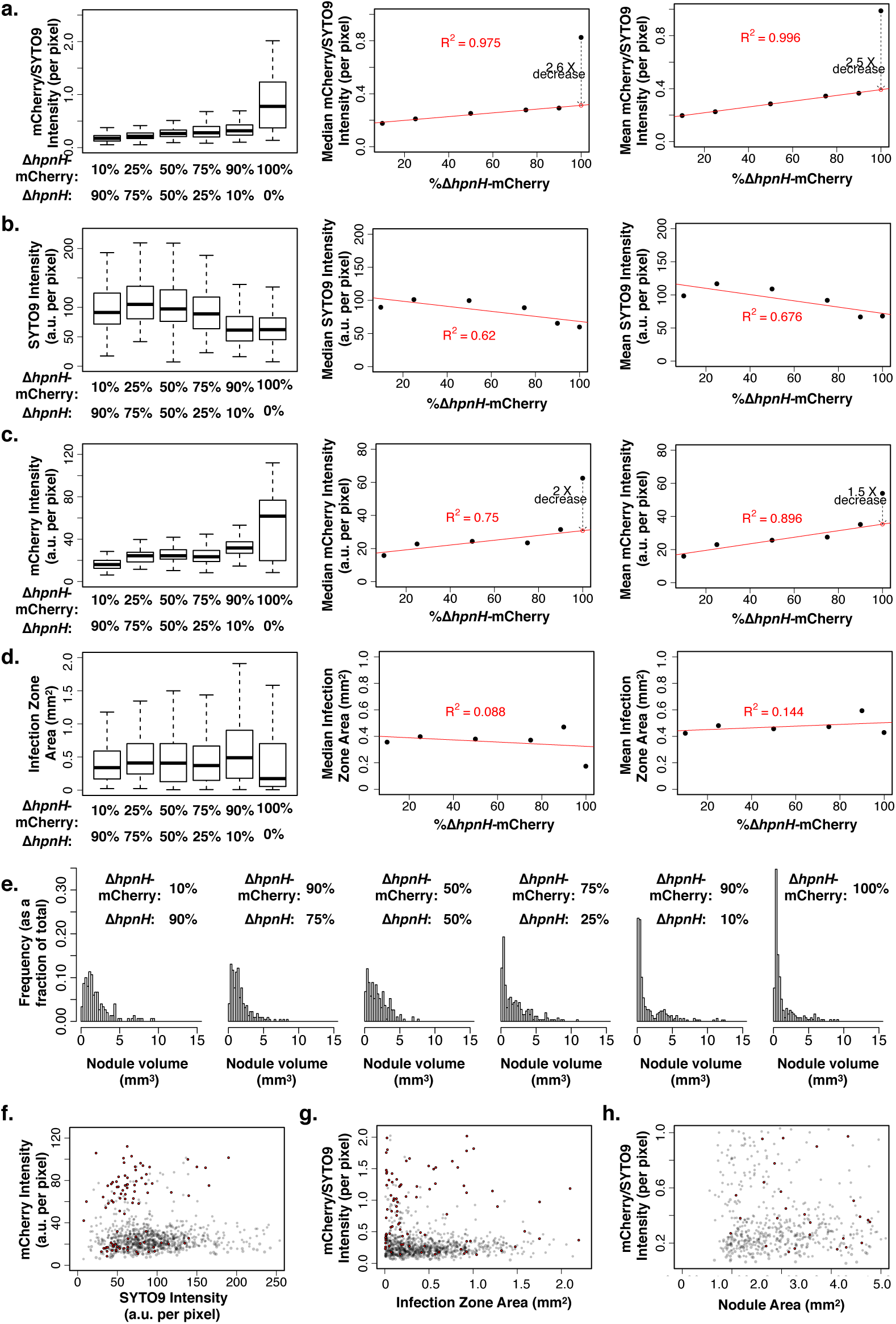
(**a**-**d**) Intensity ratio of YFP to propidium iodide (PI) (**a**), PI intensity (**b**), and YFP intensity (**c**) per pixel within infection zones of nodules co-inoculated with WT and WT-YFP strains. (**d**) Cross-sectional area of infection zones of nodules co-inoculated with WT and WT-YFP strains. For (**a**-**d**), N = 141, 95, 134, 147, 133, and 167 nodules for 10%, 25%, 50%, 75%, 90% and 100% WT-YFP strain mixtures, respectively, which were sectioned and fixed between 40-45 dpi. (**e**) Nodule volume distributions from plants co-inoculated with WT and WT-YFP strains at 40 dpi. Sample sizes are N = 183, 116, 161, 172, 232, and 248 nodules pooled from N = 8, 7, 8, 8, 8, and 8 plants for the 10%, 25%, 50%, 75% and 90% WT-YFP strain mixtures, respectively. (**f**) Scatter plots of PI vs. YFP intensities per pixel within infection zones of nodules co-inoculated with WT and WT-YFP strains. (**g**-**h**) Scatter plots of YFP/PI intensity ratios per pixel in infection zones vs. infection zone (**g**) and nodule (**h**) cross-section areas for nodules co-inoculated with WT and WT-YFP strains. Scatter plots contain data pooled from all ratios.

**Figure S18.**
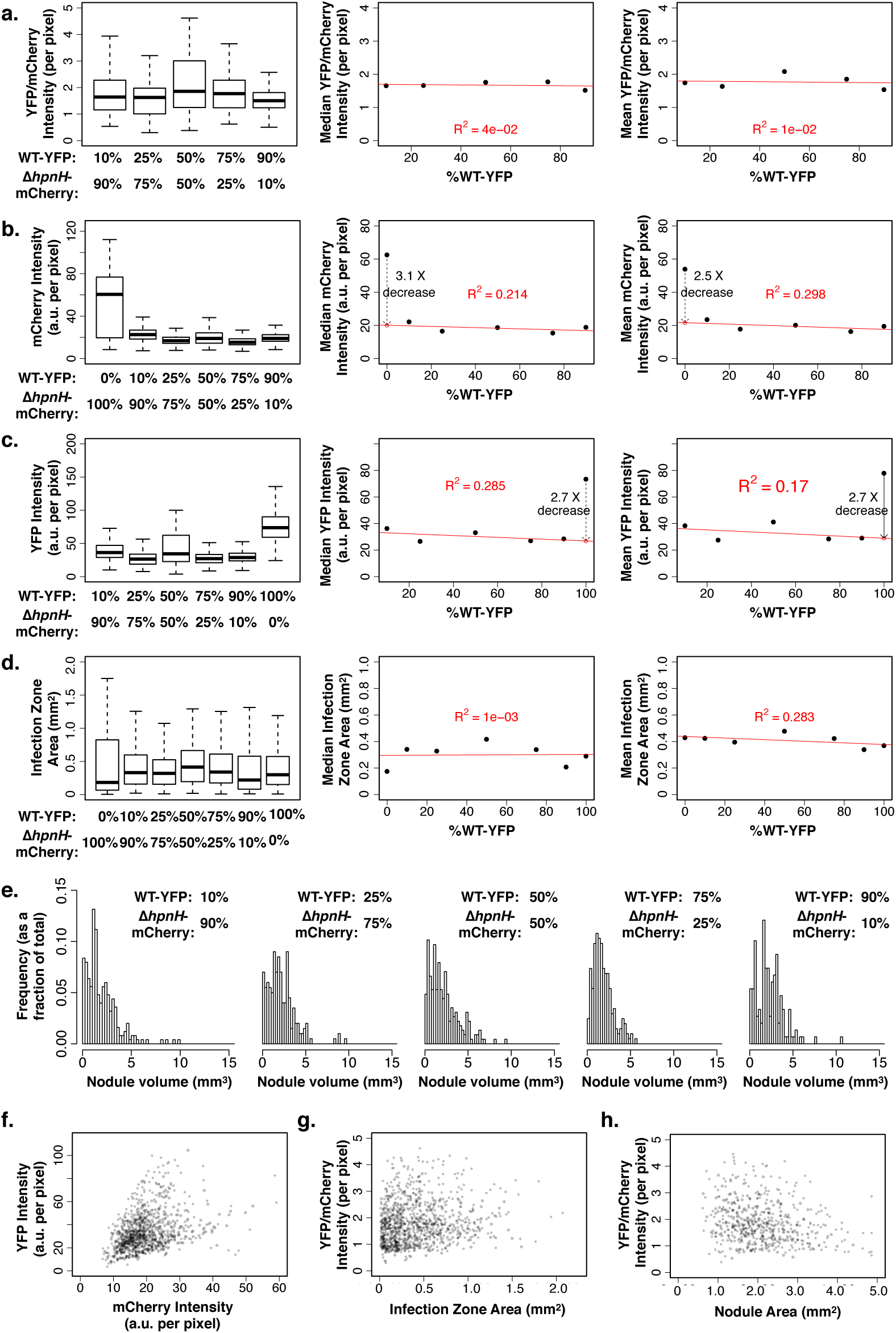
(**a**-**d**) Intensity ratio of mCherry to SYTO9 (**a**), SYTO9 intensity (**b**), and mCherry intensity (**c**) per pixel within infection zones of nodules co-inoculated with Δ*hpnH*-mCherry and Δ*hpnH* strains. (**d**) Cross-sectional area of infection zones of nodules co-inoculated with Δ*hpnH*-mCherry and Δ*hpnH* strains. For (**a**-**d**), N = 117, 107, 128, 137, 103 and 50 nodules for 10%, 25%, 50%, 75%, 90% and 100% Δ*hpnH*-mCherry strain mixtures, respectively, which were sectioned and fixed between 50-55 dpi. (**e**) Nodule volume distributions from plants co-inoculated with Δ*hpnH*-mCherry and Δ*hpnH* strains at 45 dpi. Sample sizes are N = 150, 222, 191, 254, 297, and 236 nodules pooled from N = 7, 7, 7, 8, 8, and 8 plants for the 10%, 25%, 50%, 75% and 90% WT-YFP strain mixtures, respectively. (**f**) Scatter plots of mCherry vs. SYTO9 intensities per pixel within infection zones of nodules co-inoculated with Δ*hpnH*-mCherry and Δ*hpnH* strains. (**g**-**h**) Scatter plots of mCherry/SYTO9 intensity ratios per pixel in infection zones vs. infection zone (**g**) and nodule (**h**) cross-section areas for nodules co-inoculated with Δ*hpnH*-mCherry and Δ*hpnH* strains. Scatter plots contain data pooled from all strain ratios.

**Figure S19.**
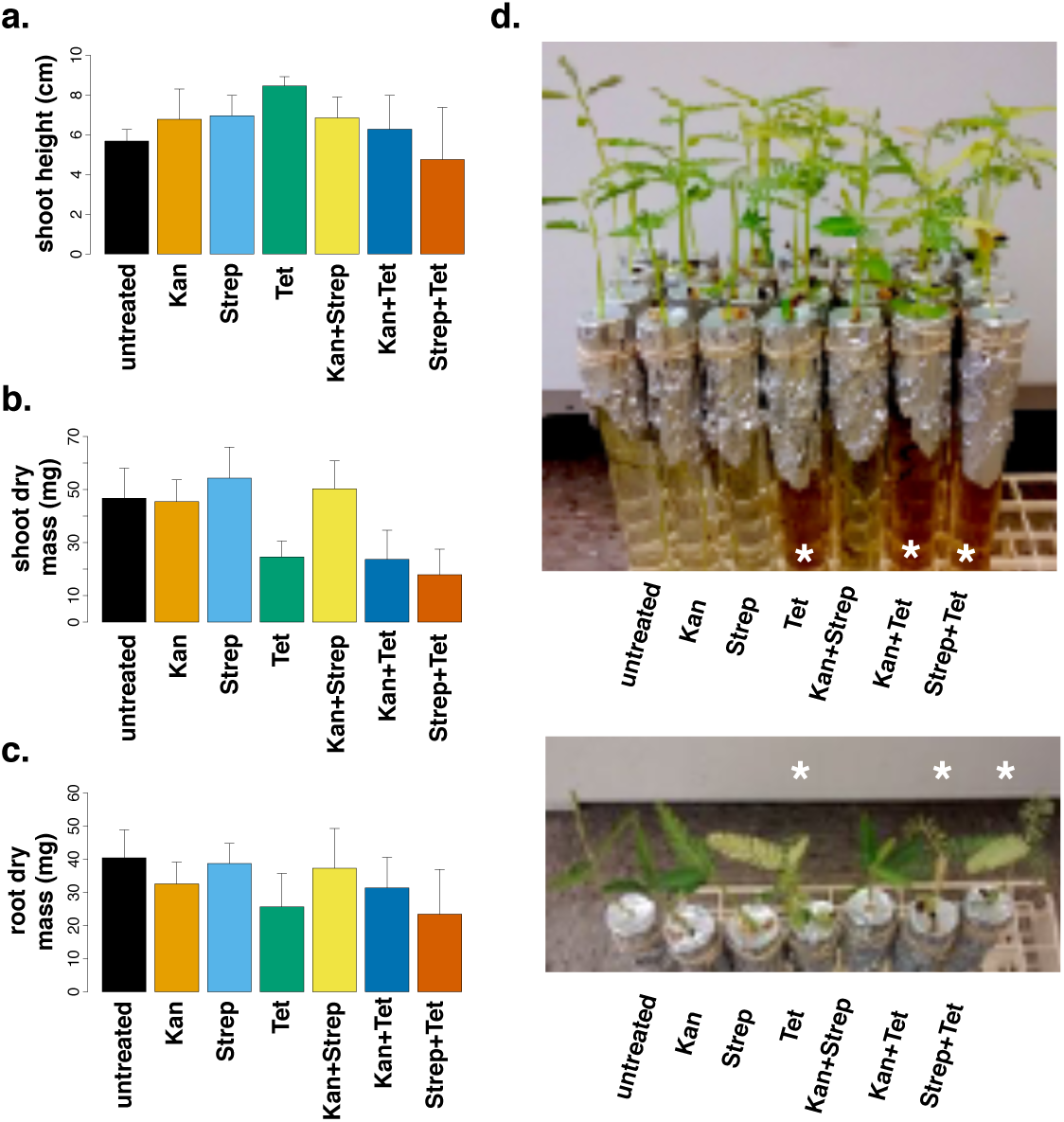
Average (**a**) shoot height, (**b**) shoot dry mass and (**c**) root dry mass for non-inoculated *A. afraspera* plants grown in BNM supplemented with kanamycin, streptomycin or tetracycline for 2 weeks under normal growth conditions. N=4 plants per condition; error bars represent one standard deviation. (**d-e**) Images of *A. afraspera* plants after 2 weeks of antibiotic treatment. Asterisks indicate plants grown in tetracycline-supplemented medium.

**Figure S20.**
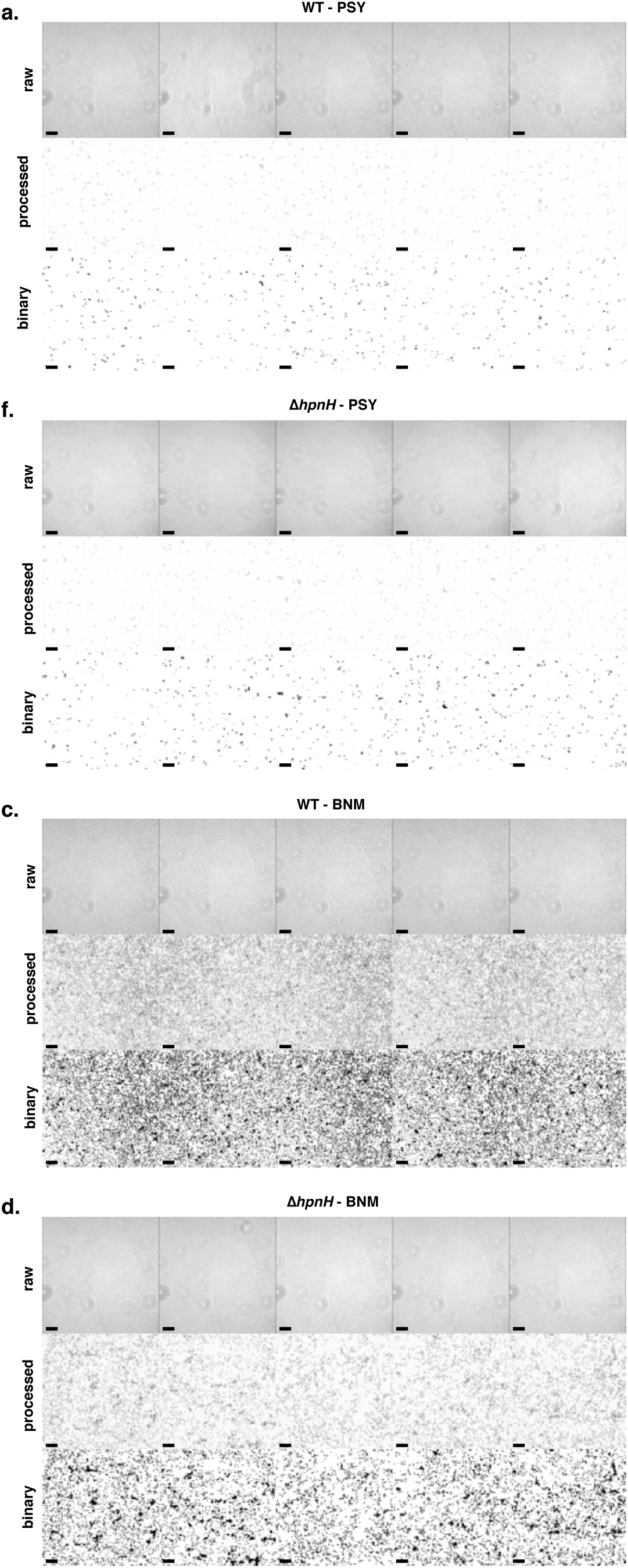
Surface attachment of wild type (**a,c**) and Δ*hpnH* (**b,d**) incubated on glass coverslips in various media. For each panel, raw phase images (top row), background-subtracted images (middle row), and binary images with cells shown in black (bottom row) are shown. Scale bars represent 20 μm.

